# Cytoskeleton remodeling induced by SMYD2 methyltransferase drives breast cancer metastasis

**DOI:** 10.1101/2023.09.18.558201

**Authors:** Alexandre G. Casanova, Gael S. Roth, Simone Hausmann, Xiaoyin Lu, Lucid Belmudes, Ekaterina Bourova-Flin, Natasha M. Flores, Ana Morales Benitez, Marcello Caporicci, Jessica Vayr, Sandrine Blanchet, Francesco Ielasi, Sophie Rousseaux, Pierre Hainaut, Or Gozani, Yohann Couté, Andres Palencia, Pawel K. Mazur, Nicolas Reynoird

## Abstract

Malignant forms of breast cancer refractory to existing therapies remain a major unmet health issue, primarily due to metastatic spread. A better understanding of the mechanisms at play will provide better insights for alternative treatments to prevent breast cancer cells dispersion. Here, we identify the lysine methyltransferase SMYD2 as a clinically actionable master regulator of breast cancer metastasis. While SMYD2 is overexpressed in aggressive breast cancers, we notice that it is not required for primary tumor growth. However, mammary-epithelium specific SMYD2 ablation increases mouse overall survival by blocking the primary tumor cells ability to metastasize. Mechanistically, we identify BCAR3 as a genuine physiological substrate of SMYD2 in breast cancer cells. BCAR3 monomethylated at lysine K334 (K334me1) is recognized by a novel methyl-binding domain present in FMNLs proteins. These actin cytoskeleton regulators are recruited at the cell edges by the SMYD2 methylation signaling and modulates lamellipodia properties. Breast cancer cells with impaired BCAR3 methylation loose migration and invasiveness capacity *in vitro* and are ineffective in promoting metastases *in vivo*. Remarkably, SMYD2 pharmacologic inhibition efficiently impairs the metastatic spread of breast cancer cells, PDX and aggressive mammary tumors from genetically engineered mice. This study provides a rationale for innovative therapeutic prevention of malignant breast cancer metastatic progression by targeting the SMYD2-BCAR3-FMNL axis.

## INTRODUCTION

Breast cancer is the most common cancer in women and the second most common cancer overall. In particular, triple-negative breast cancer (TNBC, negative for estrogen receptor ER, progesterone receptor PR and human epidermal growth factor receptor 2 HER2) and basal-like breast cancer (mostly negative for ER, PR and HER2 with expression of basal markers), are the most aggressive subtypes of breast cancer. Both have limited therapeutic options and high susceptibility to recurrence and metastases development^1, 2^. Ultimately, a vast majority of breast cancer-related deaths are due to metastatic spread to distant organs, such as lungs and bones^2, 3^. Therefore, it is important to provide alternative options preventing the development of metastases while treating primary tumors. In the course of a multi-step metastatic cascade, cancer cells acquire the ability to migrate and invade surrounding tissues^4^. Molecular mechanisms promoting invasive phenotype remain incomplete, and identifying critical molecular drivers is an urgent need to control malignant breast cancer progression^5, 6^.

Posttranslational modifications (PTMs) of proteins, the major mechanism of protein function regulation, play important roles in modulating a variety of cellular physiological and pathological processes, including human cancer hallmarks^7^. Among numerous post-translational modifications, the covalent addition of a methyl moiety on lysine residues by lysine methyltransferases (KMTs) is of particular interest. Indeed, lysine methylation signaling on both histones and non-histones proteins recently emerged as critical in normal tissue homeostasis and etiology of multiple diseases, including breast cancer^8–11^. Because of the great therapeutic potential harbored by lysine methylation due to its high specificity and reversibility, the characterization of relevant lysine methylation signaling in cancer processes may provide new therapeutic options to treat or prevent malignant cancer progression.

In this study, we utilize high-resolution single-cell RNA-seq from a cohort of breast cancer patients^12^ to decipher the metastatic potential of malignant cells associated with high expression of known active KMTs. Our analysis identifies SMYD2 (SET and MYND Domain Containing 2) as the top candidate, a lysine methyltransferase overexpressed in various cancers and previously linked to cancer pathogenesis^13–15^. Using a genetically engineered mouse model of mammary cancer, we characterize SMYD2 as a critical regulator of metastasis *in vivo*. In-depth characterization of the functional signaling at play shows that BCAR3 (Breast Cancer Anti-estrogen Resistance 3) is a genuine new SMYD2 substrate and that its mono-methylation at lysine 334 (K334me1) acts as a docking site for a new family of methyl-binding proteins, the Formin-Like proteins (FMNLs^16^). BCAR3 belongs to the NSP (novel SH2-containing protein) family, and interacts with p130/CAS at dynamic cellular adhesions where it elicits actin cytoskeleton remodeling^17–19^. FMNLs contain a synergic actin polymerization activity to boost cell migration and invasion, notably at the leading edge of motile cells in a structure called lamellipodia^20–22^. How BCAR3 and FMNLs are regulated and their functions in cancer remained unclear, and we observe that methylation of BCAR3 recruits FMNLs to increase lamellipodia fitness. Remarkably, genetic or pharmacological inhibition of SMYD2 abrogates the motility and metastatic spread of breast cancer cell lines and prevents metastatic dissemination *in vivo* in autochthonous mammary cancer models and patient-derived xenografts. Together, our data support a model in which the SMYD2-BCAR3K334me1-FMNLs axis is exploited in malignant breast cancer progression to increase lamellipodia dynamics and to promote cell motility and metastatic spreading. Our study suggests that targeting this pathway is a compelling opportunity to prevent breast cancer metastasis.

## RESULTS

### SMYD2 regulates breast cancer metastasis

Breast cancers are complex cellular ecosystems and bulk sampling methods may mask pertinent information because of the high heterogeneity of cancer tissue. To identify key lysine methyltransferases regulating breast cancer metastasis, we utilized a single-cell RNA-seq dataset of a large cohort of breast cancer patient samples representing all major clinical subtypes^12^. Restricting our analysis to neoplastic cells-only, classified in low or high metastasis potential populations based on metastasis genes signature^23^, we identified SMYD2 and EZH2 as the top two lysine methyltransferases significantly enriched in pro-metastatic cells (Fig. 1a). While EZH2 has previously been linked to breast cancer metastasis^24–26^, the participation of SMYD2 has never been demonstrated. Further bioinformatic analyses showed that *SMYD2* expression, but not *EZH2*, is elevated in breast cancer metastases compared to primary tumors (Fig. 1b and Extended Data Fig. 1a). Interestingly, while *SMYD2* is not a prognostic marker for survival of non-metastatic patients, high *SMYD2* – but not high *EZH2* – level significantly correlates with poor survival of patients diagnosed with metastatic breast cancer (Fig. 1c and Extended Data Fig. 1b-c). In addition, *SMYD2* expression is higher in aggressive TNBC and Basal-like breast cancers, two subtypes more prone to metastasis development compared to other breast cancers (Extended Data Fig. 1d). Finally, gene ontology (GO) analysis of genes which expression positively correlates with *SMYD2* upregulation in TNBC revealed an enrichment of cytoskeleton reorganization through microtubule-related programs, suggesting a potential function of SMYD2 in metastatic spread through cell motility (Extended Data Fig. 1e and Supplementary Table 1).

**Fig. 1:**
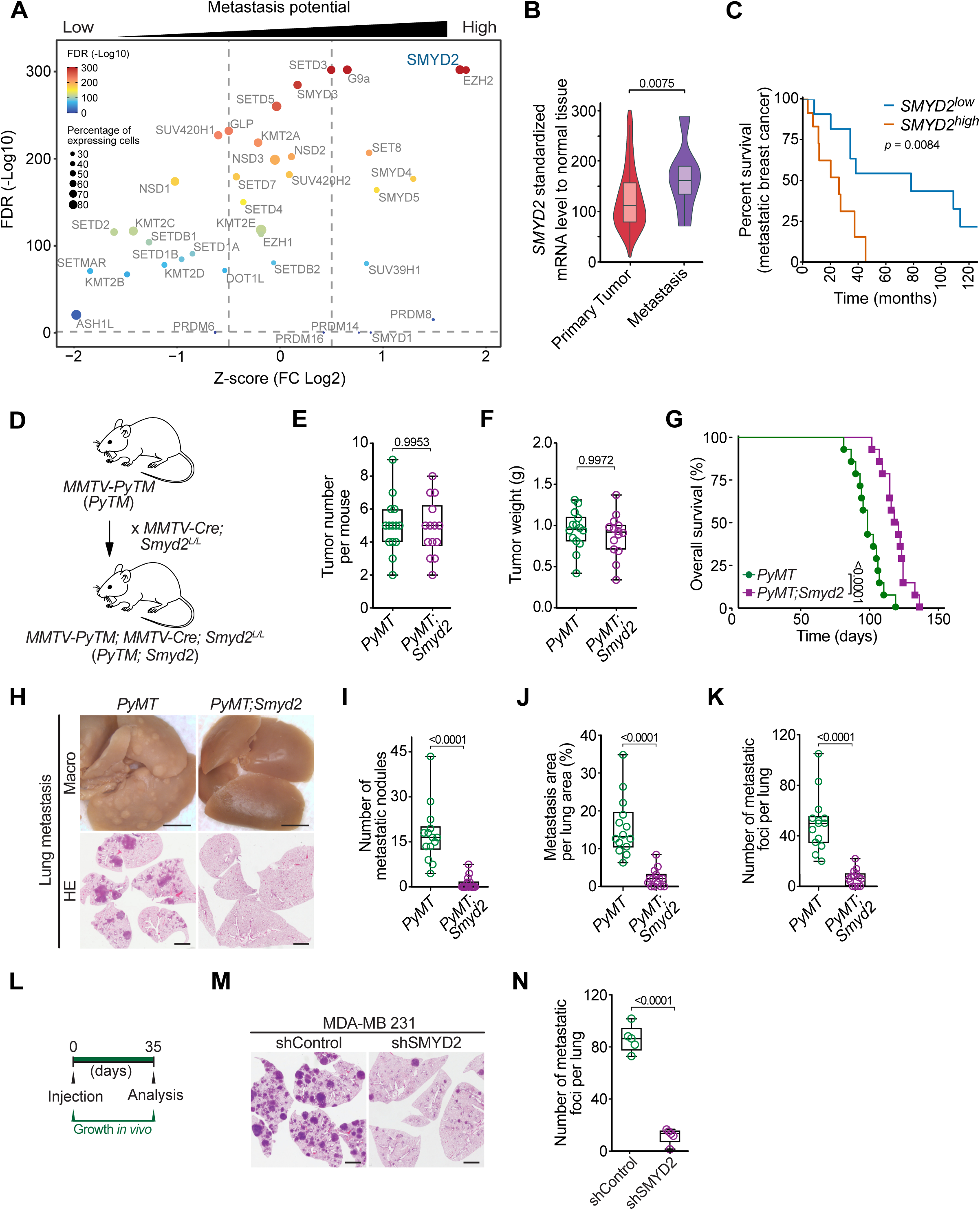
SMYD2 regulates breast cancer metastasis. **a**, Identification of lysine methyltransferases (KMTs) associated with breast cancer metastasis potential, differentially expressed in scRNA-seq of human primary breast tumors between metastasis-potential-high and -low groups. The analysis was performed using the breast cancer single-cell RNA-Seq dataset GSE176078 containing 130,246 single-cells from 26 primary pre-treatment tumors^19^. Vertical dotted lines indicate scaled Log_2_ fold changes (Z-scores) of 0.5 or::J−0.5. Adjusted *P*-values were calculated by Wilcoxon rank-sum test with Bonferroni’s correction. The color of the dot represents the adjusted *P*-value. The size of the dot represents the percentage of cells expressing the indicated gene. **b**, Violin plots of SMYD2 expression levels in primary tumor and metastases samples from breast invasive carcinoma RNA-Seq data analyzed with TNMplot^48^. *P*-values was calculated by Kruskal-Wallis test with Dunn test for multiple comparisons. **c**, Analysis of the correlation between SMYD2 expression levels and survival in a cohort of patients diagnosed with breast cancer metastatic disease from the TCGA RNAseq dataset. The SMYD2^low^ and SMYD2^high^ groups were set to the median expression of SMYD2. *P*-value were calculated by log-rank test. **d**, Schematic of *PyMT* and *PyMT;Smyd2* breast cancer mouse models generation.SMYD2 expression is elevated in breast cancer malignant progression. **e**-**f**, Quantification of tumor number (**e**) and tumor weight (**f**) in *PyMT* and *PyMT;Smyd2* mice (n = 14 mice for each experimental group). *P*-value calculated by two-tailed unpaired t test. **g**, Kaplan-Meier survival curves of *PyMT* (med. survival: 94 days, n = 14) and *PyMT;Smyd2* (med. survival: 115 days, n = 14) mice. *P*-value were calculated by log-rank test. **h**, Representative bright-field imaging and HE staining of metastatic foci in the lungs of *PyMT* and *PyMT;Smyd2* mutant mice at the endpoint. Representative of n = 14 mice for each experimental group. Scale bars, 3 mm. **i**-**k**, Quantification of number of macroscopic metastatic nodules (**i**), number of metastatic foci in the lung (**j**) and relative metastasis area (**k**) in *PyMT* and *PyMT;Smyd2* mutant mice at the endpoint. *P-*value were calculated by two-tailed unpaired t-test. **l**, Schematic of experimental design to assess the metastatic ability of SMYD2 depleted and control MDA-MB-231 cells by intravenous transplantation into recipient NSG mice. **m**-**n**, Representative HE staining (**m**) and quantification (**n**) of metastatic foci in the lungs of NSG mice injected with MDA-MB-231 cells with SMYD2 depletion. Representative of n = 5 mice for each experimental group. *P-*value were calculated by two-tailed unpaired t test. Scale bars, 3 mm. In all box plots: the center line indicates the median, the box marks the 75^th^ and 25^th^ percentiles and whiskers: min. to max. values.

Altogether, these data suggested that SMYD2 participates in breast cancer metastasis development, and to directly investigate this possibility, we utilized the *MMTV-PyMT* mouse model which develops spontaneous mammary tumors that recapitulate the tumor stages, pathology, metastasis, and biomarkers of patients with metastatic breast cancer^27^. We generated mouse models with mammary epithelium-specific SMYD2 depletion by crossing *Smyd2^L/L^* conditional knockout mice (Extended Data Fig. 1f) with *MMTV-Cre* strain. *Smyd2* gene deletion in the mouse mammary gland displayed no apparent defect in organ development and function. *MMTV-Cre;Smyd2^L/L^* mice were then interbred with *MMTV-PyMT* animals to generate *MMTV-PyMT*; *MMTV-Cre;Smyd2^L/L^* (hereafter referred to as *PyMT;Smyd2*) mutant and *MMTV-PyMT*; *MMTV-Cre* (*PyMT*) control mice (all in the FVB mouse strain, Fig. 1d and Extended Data Fig. 1g). PyMT-driven mammary tumorigenesis was examined in virgin females in the presence or absence of endogenous SMYD2.No significant difference was found in primary tumor number and weight, and both analyzed groups of mice developed extensive mammary tumor burden by 12 weeks of age (Fig. 1e-f). Consistently, no significant differences in tumor histology, proliferation, and apoptosis markers were visible between *PyMT;Smyd2* and *PyMT* tumors at 6 and 12 weeks (Extended Data Fig. 1h-j). However, we found a significant increase in *PyMT;Smyd2* mice overall survival compared to *PyMT* control animals ((24%), Fig. 1g), and the earlier morbidity observed in *PyMT* control mice was partly caused by labored breathing warranting humane euthanasia, a symptom consistent with previous observations of the *PyMT* model propensity to develop extensive lung metastasis^28^. As expected, gross examination of the lungs revealed a significant metastatic tumor burden in *PyMT* control mice, while *PyMT;Smyd2* mice had fewer visible metastatic nodules in the lungs (Fig. 1h). Detailed analyses of lung histology demonstrated a nearly 8-fold reduction in the overall number, and size of metastatic foci in *PyMT;Smyd2* vs. *PyMT* control (Fig. 1i-k), and we observed elevated SMYD2 expression in *PyMT* tumor biopsies obtained from lung metastasis vs. primary site (Extended Data Fig. 1k). To corroborate the metastasis-promoting function of SMYD2 in human breast cancer, control and SMYD2-depleted malignant MDA-MB-231 breast cancer cells were intravenously inoculated into immunocompromised mice. Histological analysis of the lungs at 35 days after injection revealed that SMYD2-depleted cells showed an 81% reduction in number of metastases relative to the control group (Fig. 1l-n).

### SMYD2 methylates BCAR3 in breast cancer cells

To decipher the mechanisms by which SMYD2 regulates breast cancer metastasis, we systematically identified potential SMYD2 methyltransferase substrates through an unbiased proteomic screen. To that end, we utilized 3xMBT (a pan-methyl binding domain) pulldowns coupled with SILAC (stable isotope labeling by amino acids in cell culture) -based quantitative proteomic approach^29, 30^. We performed a methylation assay by incubating labeled cell extracts with either recombinant wildtype SMYD2 or catalytic dead SMYD2^F184A^ mutant^31^ and compared the methylome from each condition after 3xMBT pulldowns. From over 400 identified methylated proteins by mass spectrometry (MS) -based quantitative proteomics, 25 were specifically enriched in cell extracts incubated with enzymatically active SMYD2 (Fig. 2a and Supplementary Table 2). Interestingly, six of these candidates were identified in a complementary proteomic analysis of SMYD2-dependent methylated proteins using an immune-based enrichment^32^: AHNAK, BCAR3, DBNL, DIAPH1, PRRC2C and RTF1. A correlation analysis between the expression of *SMYD2* and candidate substrates in metastatic vs. non-metastatic breast cancer identified BCAR3 as the strongest positive hit in metastatic breast cancer (Fig. 2b and Supplementary Table 3). Interestingly, BCAR3 has already been associated with breast cancer pathogenesis, however the specific mechanisms regulating BCAR3 functions remained elusive^19, 33, 34^. We confirmed the capacity of SMYD2 to methylate BCAR3 *in vitro* using recombinant proteins with *S*-adenosyl-methionine as the methyl donor (Extended Data Fig. 2a). Next, we observed by liquid chromatography-tandem mass spectrometry (LC-MS/MS) the mono-methylation of BCAR3 at lysine 334 (K334me1), which we verified by mutagenesis as the single site of methylation catalyzed by SMYD2 (Fig. 2c-d).

**Fig. 2:**
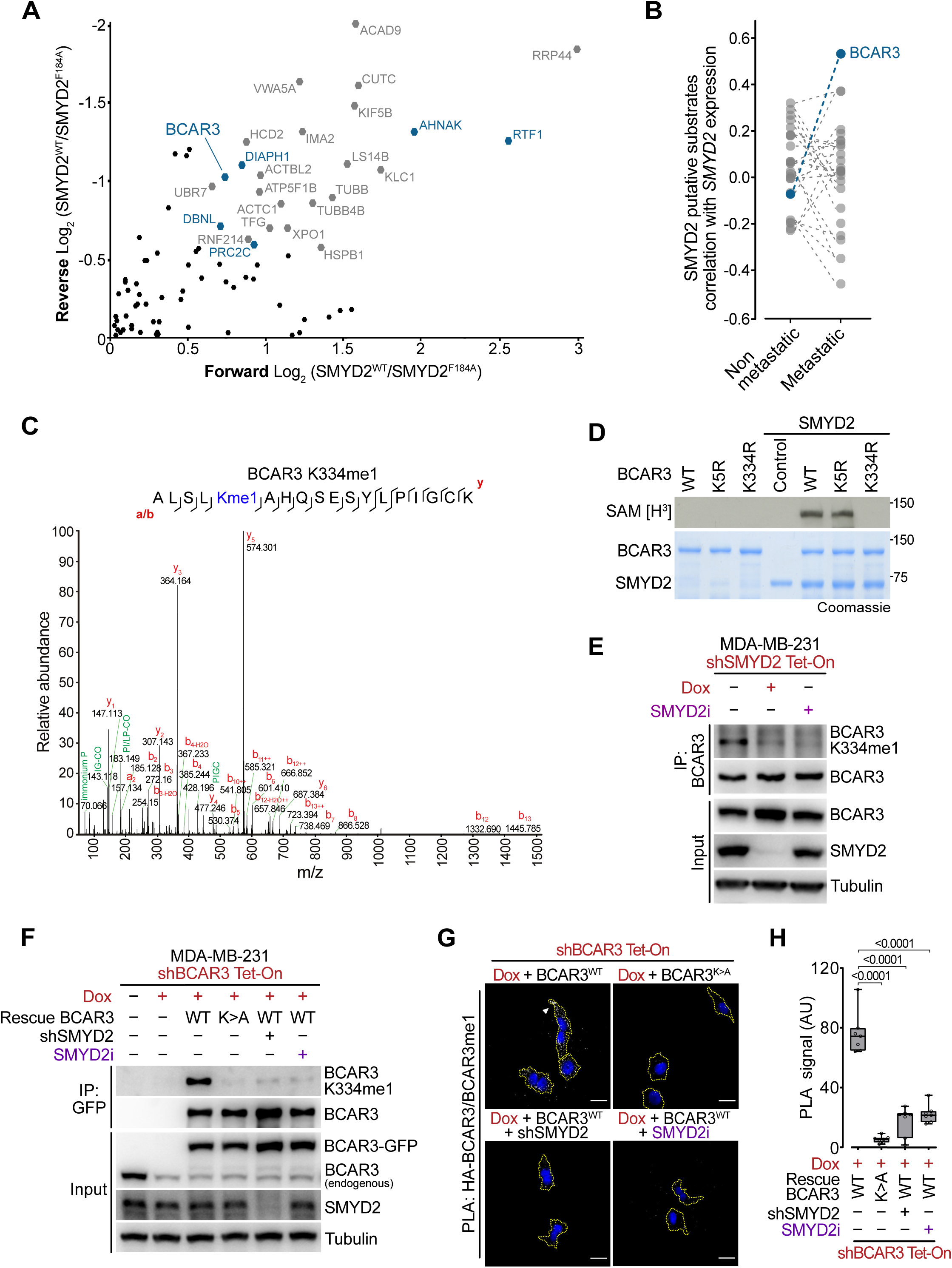
SMYD2 methylates BCAR3 in breast cancer cells. **a**, SILAC-based quantitative proteomic identification of methylome changes in SMYD2 activity proficient and deficient cell extracts. Scatter plot of proteins identified by mass spectrometry in pan-methyl-protein pulldowns (3xMBT). The X-axis shows the Log_2_ ratio of methylated proteins in the presence of active *vs* inactive SMYD2 (Forward). The Y-axis shows the Log_2_ ratio of a label-swap replicate experiment (Reverse). Potential substrates identified only in swapped replicates with a Log_2_ ratio >0.5 are shown. Candidates previously identified^26^ are labeled in blue. **b**, *BCAR3* expression significantly correlates with *SMYD2* expression in metastatic but not in non-metastatic breast cancer samples. Spearman correlation coefficients calculated for expression of genes encoding identified putative SMYD2 substrates and *SMYD2* in indicated patient groups are shown. **c**, Tandem mass spectrometry (MS/MS) spectrum identifying monomethylated K334 present on BCAR3 after *in vitro* SMYD2 methylation. Deuterated S-adenosyl-l-methionine was used as a methyl donor. **d**, Autoradiogram of a radiolabeled methylation assay using recombinant SMYD2 and full-length recombinant wildtype (WT) or point mutants K5R and K334R BCAR3 (Top panel). Bottom panel, Coomassie stain of proteins in the reaction. **e**, Immunodetection of endogenous BCAR3 K334me1 levels and upon genetic or pharmacologic SMYD2 repression in MDA-MB-157 cell line. Tubulin is shown as a loading control. **f**, Immunoblot analysis with the indicated antibodies of whole cell lysate and immunoprecipitated (IP) proteins from MDA-MB-231 cells with doxycycline-induced (Dox) shRNA depletion of BCAR3 and rescue complementation with GFP-tagged wildtype (WT) or K334A (K>A) BCAR3, in combination with SMYD2 depletion (shSMYD2) or pharmacologic repression (SMYD2i). Tubulin is shown as a loading control. **g**-**h** Representative images (**g**) and signal quantification (**h**) of proximity ligation assay (PLA) detecting methylated BCAR3 by coupling antibodies against HA-tagged total BCAR3 and BCAR3 K334me1 in MDA-MB-231 cells carrying indicated modifications. Dotted yellow lines represent cells periphery and arrow identifies enriched signal at cell edge. *P*-values were calculated by ANOVA with Tukey’s testing for multiple comparisons. Scale bars, 10μm.

To investigate BCAR3 methylation in cells, we raised a methyl-specific antibody against BCAR3 K334me1, that proved to be highly specific for BCAR3 K334me1 (Extended Data Fig. 2b). We confirmed that the methyl-BCAR3 antibody detects full-length recombinant methylated BCAR3, but not the unmethylated, K334R mutant, nor wildtype BCAR3 incubated with the selective SMYD2 catalytic inhibitor BAY-598^35^ (Extended Data Fig. 2c-d). Finally, we detected BCAR3 methylation using 293T cell lysates expressing ectopic SMYD2 and BCAR3 (Extended Data Fig. 2e). We then assessed the endogenous levels of BCAR3 in a panel of breast cancer cell lines BCAR3 (Extended Data Fig. 2f). We found that BCAR3 expression was elevated in metastatic cell lines MDA-MB-231 and MDA-MB-157 (basal-like/TNBC) compared to less invasive MDA-MB-468 (basal-like/TNBC) and MDA-MB-453 (HER2+ subtype), and BCAR3 was low-to-absent in non-invasive MCF-7 (luminal) cell lines. In addition, SMYD2 level was relatively similar in all breast cancer cell lines tested. Corresponding with elevated BCAR3 and SMYD2 levels, we detected a robust endogenous BCAR3 methylation signal using the specific BCAR3 K334me1 antibody in the metastatic breast cancer lines (Fig. 2e and Extended Data Fig. g-h). Importantly, the BCAR3 methylation was depleted upon genetic (inducible shRNA) or pharmacologic (BAY-598 inhibitor) suppression of SMYD2. In contrast, we did not observe BCAR3 methylation in less or non-invasive breast cancer cells (Extended Data Fig.2i).

Next, we generated MDA-MB-231 cells with shRNA-mediated depletion of endogenous SMYD2 or doxycycline-inducible depletion of endogenous BCAR3 (Dox) and ectopic expression of wildtype or methyl-mutant (K334A) BCAR3 or corresponding control. Using these engineered cells to modulate actors of the SMYD2-BCAR3 methylation signaling, we confirmed that restored wildtype BCAR3 but not K334A mutant is methylated by SMYD2 in these cell lines, and that the K334 methylation is lost upon genetic or pharmacologic repression of SMYD2 (Fig. 2f). In agreement with our initial observations in the mouse model (Extended Data Fig. 1f-h), SMYD2 ablation had no impact on MDA-MB-231 proliferation (Extended Data Fig. 2j). Of note, BCAR3 depletion slightly decreased cell proliferation as previously observed^36^, but independently of its methylation, as the K334A BCAR3 mutant was able to rescue proliferation defect. Finally, we used proximity ligation assay (PLA) with antibodies detecting total and K334me1 BCAR3 to assure specificity of methylated BCAR3 signal, and observed *in situ* methylated BCAR3 in the cytoplasm with a diffused enrichment at the cell leading edge (Fig. 2g). As expected, BCAR3 K334me1 signal was lost upon genetic or pharmacological suppression of SMYD2 or expression of the methyl-mutant BCAR3 K334A (Fig. 2g-h).

### BCAR3 methylation promotes cell migration and invasiveness

Our observations suggested a potential role for SMYD2-mediated BCAR3 methylation at K334 in malignant breast cancer pathogenesis. To assess the potential role of BCAR3 K334me1 in the regulation of cancer cell phenotypes, we performed genetic complementation experiments using previously described engineered MDA-MB-231 cell lines. We quantified single-cell migration velocity with time-lapse live microscopy and found that SMYD2 or BCAR3 depletion significantly impaired individual cell motion (Fig. 3a-b). Interestingly, complementation with wildtype, but not BCAR3 harboring a K334A substitution, could fully restore MDA-MB-231 cancer cell migration velocity. This led us to speculate that the SMYD2-BCAR3 methylation signaling is exploited by cancer cells to activate a pro-migratory phenotype. To test this hypothesis, we ectopically expressed wildtype or K334A BCAR3 in non-invasive MCF-7 breast cancer cells characterized by the absence of BCAR3 methylation (Extended Data Fig. 2f and 3a). Of note, while we also detected modest proliferation changes upon BCAR3 expression in these cells, those were independent of BCAR3 methylation status (Extended Data Fig.3b). Remarkably, we observed a significant increase in MCF-7 cell migration with wildtype methylatable BCAR3 compared to control or K334A BCAR3 (Extended Data Fig. 3c-d).

**Fig. 3:**
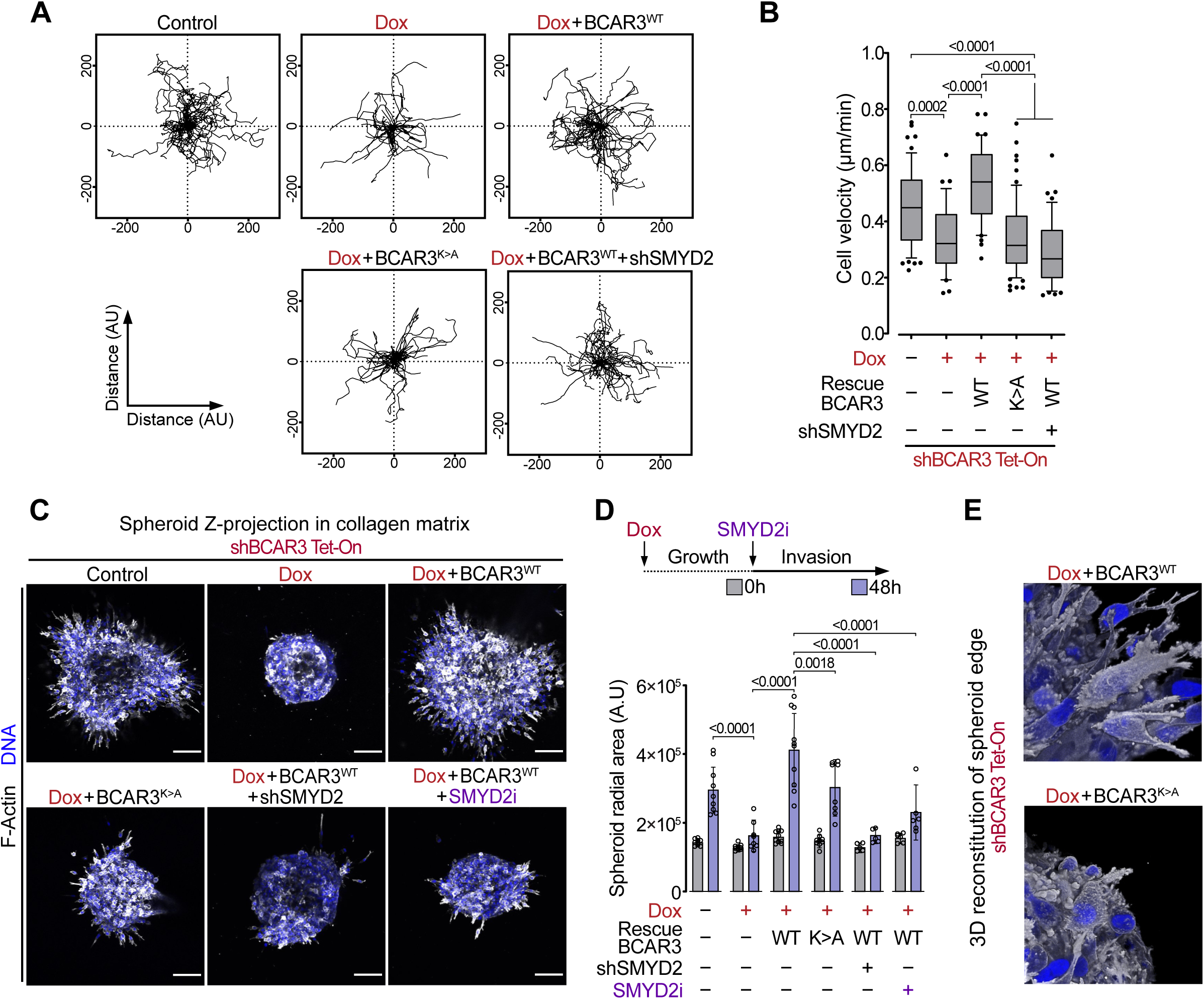
BCAR3 methylation promotes cell migration and invasiveness. **a**-**b**, Representative single-cell migration tracks monitored by time-lapse live microscopy (**a**) and quantification of single-cell migration velocity (**b**) of MDA-MB-231 cells with indicated engineering. Boxes: 25^th^ to 75^th^ percentile, whiskers: 10 to 90 percentile, center line: median. *P*-values were calculated by ANOVA with Tukey’s testing for multiple comparisons. **c**-**d**, Representative images of three-dimensional collagen I matrix invasion (**c**) and spreading quantification (**d**) of basement membrane enclosed MDA-MB-231 with indicated engineering. *P*-values were calculated by ANOVA with Tukey’s testing for multiple comparisons. Scale bars, 50μm. **e**, 3D image reconstruction of spheroid edge depicting engineered MDA-MB-231 cells protrusions, related to (**c**). Scale bars represent 10µm. In all box plots, the center line indicates the median, the box marks the 75^th^ and 25^th^ percentiles and the whiskers indicate 10 to 90 percentile values.

A critical step in metastasis is epithelial invasion, during which cancer cells pass the basement membrane and migrate through the underlying collagen-rich stromal extracellular matrix thanks to a dynamic orchestration of cell motility and matrix anchorage processes. To assess the role of the SMYD2-BCAR3 methylation signaling in invasion, we used a three-dimensional collagen matrix model in which spheroids of MDA-MB-231 enclosed by basement membrane Matrigel were embedded in thick collagen I hydrogel. We found that BCAR3 depletion significantly impairs MDA-MB-231 cells matrix invasion, a phenotype rescued by complementation with wildtype BCAR3, but not with the K334A mutant (Fig. 3c-d). In agreement with these results, genetic and pharmacologic repression of SMYD2 phenocopied BCAR3 ablation and BCAR3 K334A mutation. Finally, detailed confocal adaptive optics microscopy analyses of cancer cells migrating from the spheroid mass showed that cells impaired for BCAR3 methylation presented protrusions defects (Fig. 3e). These protrusions are a constitutive feature of highly metastatic cells providing the primary mechanical impetus for physiological motility, enabling cell spreading via mechanical and proteolytic matrix remodeling^5, 21^. Therefore, our data suggested that BCAR3 methylation participates in malignant cell protrusions remodeling.

### FMNLs are BCAR3 methyl-specific interactors

Lysine methylation predominantly regulates protein-protein interactions^37^. Therefore, to identify K334 methylation-sensitive binding partners of BCAR3, we performed a comprehensive SILAC-based quantitative proteomic screen with MDA-MB-231 cells extracts to isolate proteins that bound differentially to BCAR3 K334me0 versus K334me1 peptides. Mass spectrometry analyses revealed three strong candidates that bind specifically to methylated BCAR3. These identified methyl-specific interactors, FMNL1, FMNL2, and FMNL3 (hereafter referred to as FMNLs), are the three members of the Formin-like protein family (Fig. 4a and supplementary Table 4). Notably, FMNLs are known to bundle and polymerize actin filaments (F-Actin) and participate in actin cytoskeleton remodeling during cell migration and metastatic progression^20, 22, 38^. We noticed that FMNL2 and FMNL3 were upregulated in metastatic breast cancer cell lines (Extended Data Fig. 4a). Next, we confirmed the interaction between BCAR3 K334me1 peptides and endogenous FMNL2/3 using MDA-MB-231 cells extract and additionally found that this interaction was direct using recombinant FMNLs (Fig. 4b-c). We showed that endogenous FMNL2 and FMNL3 co-immunoprecipitated with endogenous methylatable BCAR3 in MDA-MB-231 cells, and this interaction was lost upon SMYD2 genetic and pharmacologic inhibition (Fig. 4d). Additionally, we made similar observations with BCAR3-depleted MDA-MB-231 cells rescued with wildtype BCAR3, while no interaction was detectable with K334A BCAR3 or repressed SMYD2 (Extended Data Fig. 4b). Next, we tracked the localization of direct BCAR3-FMNL3 interaction by PLA and observed that FMNL3 interacted *in situ* with methyl- proficient wildtype BCAR3 but not with the methyl-deficient BCAR3 K334A nor upon SMYD2 inhibition (Fig. 4e-f). Importantly, we noted that the BCAR3-FMNL3 interaction localized predominantly at the cell edges where FLMN3 could promote migrating force through cytoskeleton remodeling.

**Fig. 4:**
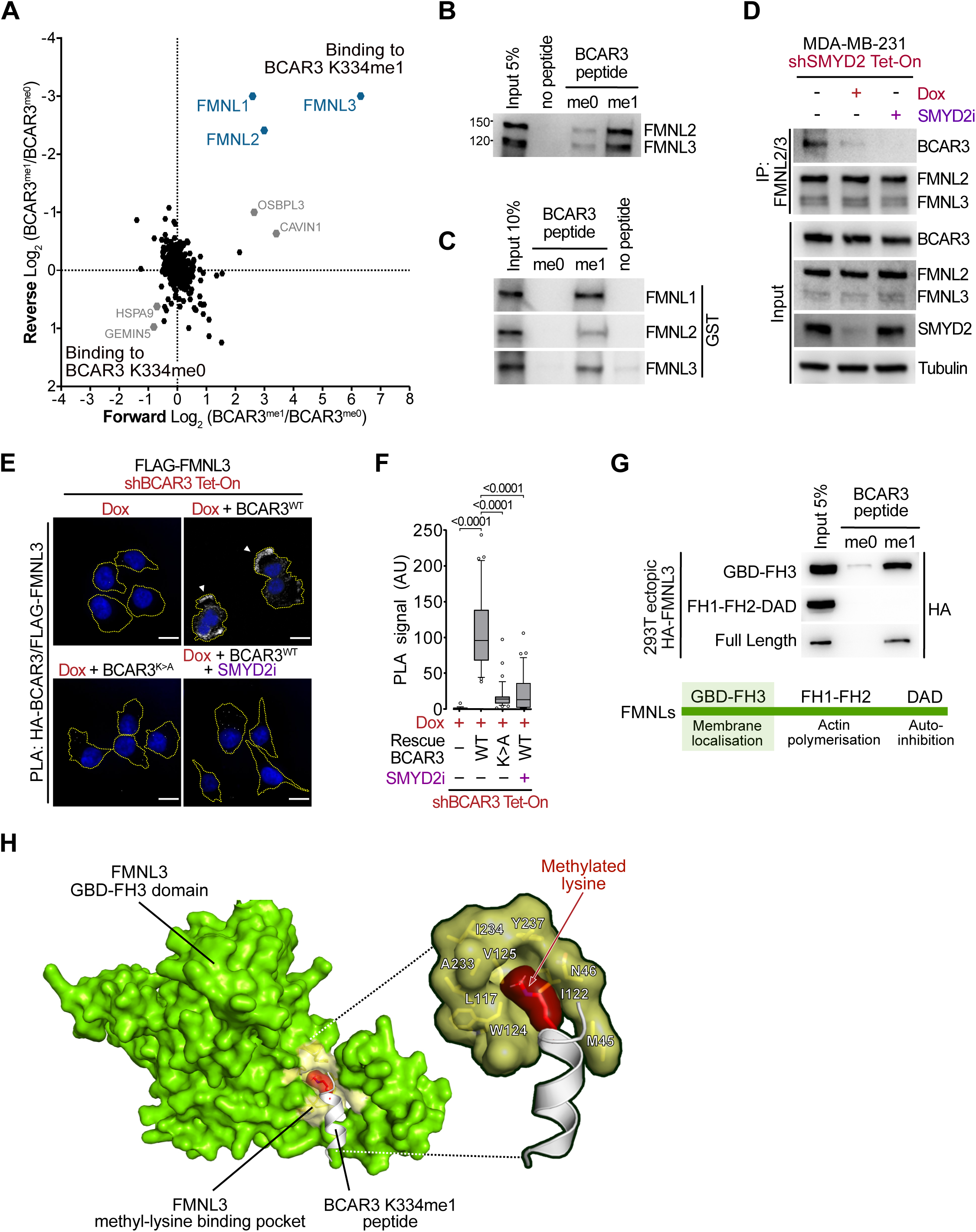
FMNLs are BCAR3 methyl-specific interactors. **a**, Identification of K334 methylation-specific BCAR3 binding partners using SILAC-based quantitative proteomics screen. A scatter plot of the log_2_ transformed protein-normalized methyl-sensitive peptide SILAC ratios. The X-axis shows the Log_2_ ratio of proteins binding to BCAR3 K334me1 versus BCAR3 K334me0 (Forward). The Y-axis shows the Log_2_ ratio of a label-swap replicate experiment (Reverse). Specific interactors of methylated BCAR3 reside in the upper right quadrant and methyl-sensitive binders identified in swapped replicates with a Log2 ratio > ±0.5 are presented. **b-c**, Immunoblot analysis with the indicated antibodies of protein pulldowns using unmethylated (me0) or K334 monomethylated (me1) BCAR3 peptides from MDA-MB-231 cell extracts (**b**) or recombinant GST-FMNLs proteins (**c**). **d**, Co-immunoprecipitation of endogenous BCAR3 after enrichment of endogenous FMNL2/3 in MDA-MB-231 cells upon genetic or pharmacologic SMYD2 repression. Tubulin is shown as a loading control. **e**-**f**, Representative images (**e**) and signal quantification (**f**) of PLA monitoring BCAR3-FMNL3 interaction *in situ*, coupling HA-BCAR3 and Flag-FMNL3 antibodies in MDA-MB-231 cells with indicated engineering. Dotted yellow lines represent cells periphery and arrows identify enriched signal at cellular edge. *P*-values were calculated by Brown-Forsythe and Welch ANOVA tests with Dunnett’s T3 testing for multiple comparisons. Scale bars, 10μm.**g**, Immunoblot analysis with the indicated antibodies of protein pulldowns using unmethylated (me0) or K334 monomethylated (me1) BCAR3 peptides from extracts of 293T cells expressing full-length, GBD-FH3 or FH1-FH2-DAD domains of FMNL3. **h**, Predicted structure model of FMNL3 GBD-FH3 domain interaction with BCAR3 K334me1 peptide, based on the available structure of FMNL2 GBD-FH3 / CDC42 GppNHp (PDBe - 4YC7). The putative methylated lysine-binding hydrophobic pocket of FMNL3 containing two aromatic residues (W124 and Y237) surrounded by aliphatic residues (L117, I122, V125, A233) is shown. In all box plots, the center line indicates the median, the box marks the 75^th^ and 25^th^ percentiles and the whiskers indicate 10 to 90 percentile values.

Intriguingly, FMNLs lack any characterized methyl-binding domain. Analysis of the domains shared by Formin proteins with publicly available structural information revealed that the N-terminal regulatory fragment comprising the GTPase-binding domain and FH3 domain (GBD-FH3) is organized as an Armadillo-repeat fold (Extended Data Fig. 4c), which shares a high structural redundancy with HEAT repeats^39^ known to interact with mono-methylated lysine 20 of H4^40^. We confirmed that the GBD-FH3 domain of FMNL3 was sufficient to specifically bind methylated BCAR3 (Fig. 4g and Extended Data Fig. 4d). The molecular structure of the GBD-FH3 domain of FMNL2 in complex with the Rho-GTPase Cdc42 was previously determined^41^. We performed a structural homology modeling of the BCAR3-FMNL3 interaction based on the high structural similarity between FMNL2 and FMNL3, and between the Cdc42 helix α3 responsible for its interaction with the GBD-FH3 domain and the sequence of the methylated BCAR3 peptide (Fig. 4h). Remarkably, our model indicated that monomethylated lysine K334 of BCAR3 is located within a well-defined hydrophobic pocket of FMNL3 containing two aromatic residues (W124 and Y237) surrounded by aliphatic residues (L117, I122, V125, A233), which defined an optimal methylated lysine-binding ‘aromatic cage’ structural motif^42^. Importantly, the six critical amino acids defining the hydrophobic pocket of FMNL3 are strictly conserved in FMNL1 and FMNL2 (Extended Data Fig. 4e-f).

### BCAR3 methylation recruits FMNLs to lamellipodia and regulates cell protrusions

FMNLs are known to facilitate the formation of protrusive cytoskeleton structures such as lamellipodia at the leading edge of migrating cells^20^. As the actin-based protrusions are confined to distinct cellular compartments, a membrane-targeting mechanism is critical for the function of FMNLs^21^. The GBD-FH3 domain was previously shown to control FMNLs localization to cell membrane either directly by N-terminal myristoylation anchoring or indirectly by binding to Rho-GTPases such as Cdc42, which opens and activates FMNLs^22, 41^. Based on our previous observations, we speculated that BCAR3 methylation may recruit and activate FMNLs effectors to promote cell migration. We performed immunocytostaining of MDA-MD-231 cells and observed a clear cytoplasmic signal for both SMYD2 and BCAR3, accumulated at cell protrusions resembling lamellipodia (Extended Data Fig. 5a-b). Remarkably, confocal microscopy confirmed that FMNL3 strongly accumulated in the cytoplasm and accumulated at lamellipodia in cells proficient for the SMYD2-BCAR3 signaling, but not when BCAR3 methyl-mutant or SMYD2 catalytic inhibitor were used (Fig. 5a-b).

**Fig. 5:**
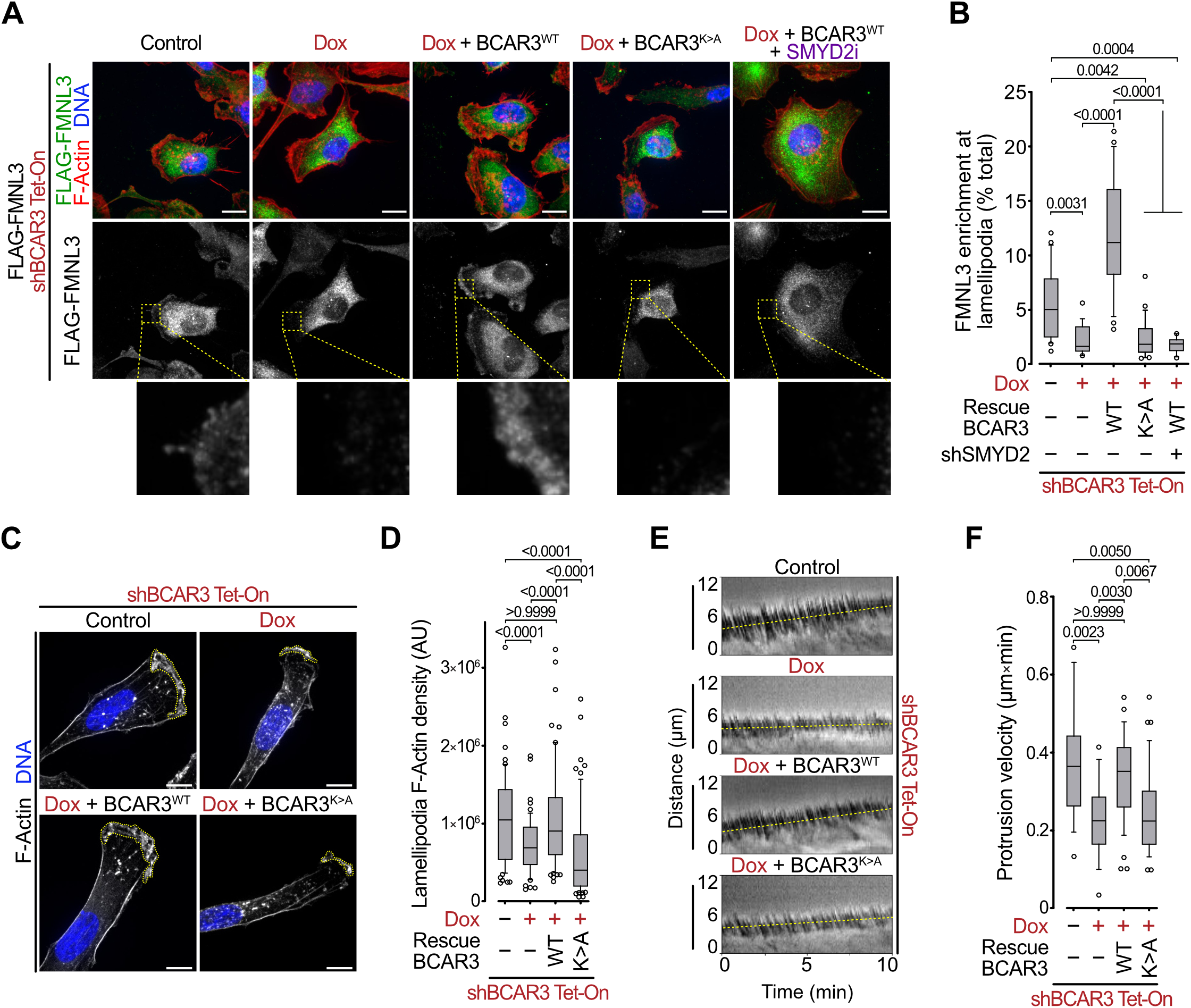
BCAR3 methylation recruits FMNLs to lamellipodia and regulates cytoskeleton protrusions. **a-b**, Representative immunofluorescence images (**a**) and signal quantification (**b**) of F-Actin (phalloidin) and Flag-FMNL3 in MDA-MB-231 cells with indicated engineering. Magnifications of the leading edge of lamellipodia protrusions are provided. *P*-values were calculated by Brown-Forsythe and Welch ANOVA tests with Dunnett’s T3 testing for multiple comparisons. Scale bars, 5μm. **c**-**d**, Representative images of lamellipodia visualized by Z-projection of F-Actin staining (**c**) and lamellipodia F-Actin density quantification (**d**) in MDA-MB-231 cells with indicated engineering. Dotted yellow lines represent lamellipodia area. *P*-values were calculated by Brown-Forsythe and Welch ANOVA tests with Dunnett’s T3 testing for multiple comparisons. Scale bars, 5μm. **e**-**f**, Representative visualization (kymograph, distance vs time) of lamellipodia protrusion rate (**e**) and quantification of lamellipodia protrusion velocity (**f**) in MDA-MB-231 cells with indicated modifications. *P*-values were calculated by Kruskal-Wallis test with Dunn’s testing for multiple comparisons. In all box plots, the center line indicates the median, the box marks the 75^th^ and 25^th^ percentiles and the whiskers indicate 10 to 90 percentile values.

FMNL2 and FMNL3 nucleate and elongate actin filaments to generate the force required for cell migration^20^. To investigate the functional importance of the SMYD2-BCAR3 methylation signaling in lamellipodia fitness, we performed high-resolution microscopy analyses in MDA-MB-231 cells. We first noted that the cells with BCAR3 depletion exhibited severe morphological changes and a significant decrease in lamellipodia size, a phenotype rescued with wildtype but not with the methyl-mutant K334A BCAR3 (Extended Data Fig. 5c-d). Quantification of F-actin in the lamellipodia cytoskeleton network revealed a significant reduction of the F-actin filaments density in cells depleted for endogenous BCAR3 or specifically impeded for BCAR3 methylation (Fig. 5c-d). To test if the SMYD2-BCAR3 signaling is sufficient to promote protrusions, we used MCF-7 cells with ectopic expression of wildtype or methyl-mutant K334A BCAR3. We found that BCAR3 methylation was able to significantly increase F-actin network density at cell protrusions, although fully polarized lamellipodia were not visible (Extended Data Fig. 5e-f). Finally, to quantify the effects of BCAR3 K334 methylation on lamellipodia velocity, we used phase contrast live microscopy to measure cell protrusion rates. Our analysis revealed a significantly decreased velocity of lamellipodium protrusions upon depletion of BCAR3, which was efficiently restored by complementation with wildtype but not K334 methyl-mutant BCAR3 (Fig. 5e-f).

### SMYD2-BCAR3-FMNLs axis drive breast cancer metastasis *in vivo*

We sought to determine the role of SMYD2-BCAR3 methylation signaling in the regulation of metastatic colonization using *in vivo* models of breast cancer. To model metastatic colonization of the lung, we intravenously inoculated highly lung-metastatic prone MDA-MB-231 breast cancer cells with depletion of endogenous SMYD2 and/or BCAR3 complemented with wildtype or methyl-mutant K334A BCAR3. These cell lines were also stably labeled with an Akaluc reporter^43^ to facilitate the quantification of lung metastasis by bioluminescence imaging (BLI). Analysis shortly after injection indicated that cells in all experimental groups became trapped in the lung capillaries, and none of the introduced genetic or pharmacological modalities impacted the number of cancer cells in the lung (Fig. 6a, day 0 and Extended Data Fig. 6a). After 35 days, we observed elevated BLI signals in the lungs of animals which received control and wildtype BCAR3 rescued cells (Fig. 6a-b, day 35). In contrast, animals inoculated with cells depleted for BCAR3 or SMYD2 or with impaired BCAR3 methylation showed significantly lower BLI signal. Additional histological analysis confirmed multiple tumor foci formed only by the cells proficient in the SMYD2-BCAR3 methylation signaling (Fig. 6a, lower panels).

**Fig. 6:**
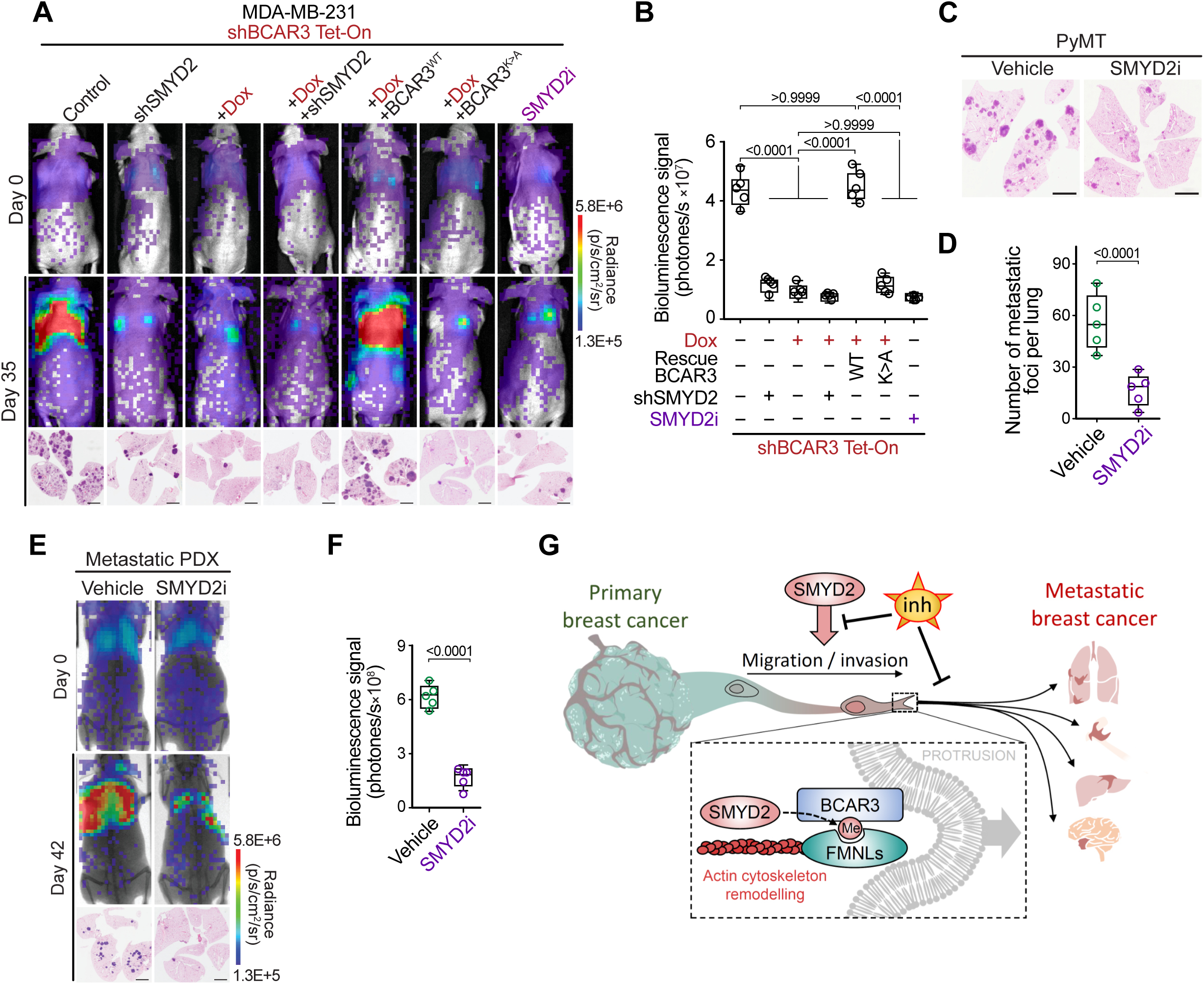
The SMYD2-BCAR3-FMNLs axis drive breast cancer metastasis. **a**, Representative bioluminescence imaging of animals at the time of intravenous transplantation (day 0) and 35 days post-injection of MDA-MB-231 cells with indicated modifications. Lower panel, representative HE staining of the lungs at day 35. Representative of n = 5 mice for each experimental group. Scale bars, 3 mm. **b**, Quantification of bioluminescence signal corresponding to metastatic cancer growth in animals as in (**a**) at day 35. Representative of n = 5 mice for each experimental group. *P*-values were calculated by ANOVA with Tukey’s testing for multiple comparisons. **c-d**, Representative HE staining (**c**) and quantification (**d**) of metastatic foci in the lungs of *PyMT* mice treated with SMYD2i inhibitor or vehicle (control) at 12 weeks of age. Representative of n = 5 mice for each experimental group. *P*-value were calculated by two-tailed unpaired t-test. **e**-**f**, Representative bioluminescence visualization (**e**) and signal quantification (**f**) of breast cancer cells obtained from patient-derived xenografts and intravenously transplanted into recipient NSG mice and treated with SMYD2i inhibitor or vehicle (control). Lower panel representative HE staining of metastatic foci in the lungs at day 42. Representative of n = 5 mice for each experimental group. Scale bars, 3 mm. *P*-value were calculated by two-tailed unpaired t-test. **g**, SMYD2 regulates breast cancer cells motility and metastasis spreading through methylation of BCAR3 and recruitment of FMNLs to protrusive membrane structures such as lamellipodia. FMNLs localization at nascent lamellipodia enables nucleation and elongation of actin filaments and generates the force required for cell migration. Hence, upregulation of the SMYD2-BCAR3-FMNLs pathway, commonly observed in the malignant breast cancer cell, promotes metastatic spread. Pharmacological inhibition of SMYD2 enzymatic activity in pre-clinical *in vivo* animal models can efficiently prevents breast cancer’s ability to metastasize. In all box plots, the center line indicates the median, the box marks the 75^th^ and 25^th^ percentiles and whiskers: min. to max. values.

To attest the pre-clinical value of targeting the SMYD2-BCAR3 pathway to prevent breast cancer dissemination, we performed *in vivo* metastasis experiment with the selective SMYD2 catalytic inhibitor BAY-598^35^. Our study revealed that pharmacological attenuation of SMYD2 activity significantly inhibited MDA-MB-231 cells metastatic spread *in vivo* (Fig. 6a-b). Concordant to previous studies showing the tolerance of healthy tissues to SMYD2i^31^, no severe side effects, including weight loss, were observed in the mice treated with SMYD2i (data not shown).

To corroborate the metastasis-promoting function of the SMYD2-BCAR3 signaling, we used the non-invasive MCF-7 cell line. In agreement with previous observations^44^, control MCF-7 cells rarely formed metastasis in mice, while cells with concomitant expression of SMYD2 and BCAR3 exhibited a significant increase (27-fold) in the number of metastatic foci in the lung (Extended Data Fig. 6b-d). Importantly, MCF-7 cells overexpressing K334A methyl-mutant BCAR3, even when combined with overexpression of SMYD2, failed to form metastases *in vivo*. In addition to the lungs, one of the most common sites of metastasis for breast cancer are bones. To test SMYD2i efficacy to prevent bone metastasis, we utilized the SUM159-M1a breast cancer subline, shown to generate robust distal metastasis, including into the bones^45^. Cells were labeled with a bioluminescence reporter and treated with SMYD2i for 5 days prior to injection to the left ventricle, to disseminate cells via the arterial circulation to the whole body. We observed the development of bone metastases in control mice, while SMYD2i treatment significantly reduced the number of metastases to distal organs (Extended Data Fig. 6e-f). Next, we sought to verify if SYMD2i can phenocopy our observations in the *PyMT* mammary cancer mouse model, in which *Smyd2* knockout largely blocked metastasis development (Fig. 1e-h). We initiated SMYD2i treatment of *PyMT* mice at the time when palpable mammary tumors were detectable and continued until control animals showed signs of morbidity (Extended Data Fig. 6g). Histological analysis revealed an over 4-fold reduction of metastatic foci in the lungs of animals treated with SMYD2i compared to the control. (Fig. 6c-d). Finally, we isolated cells from patient-derived xenograft (PDX) originating from aggressive basal-like breast cancer and performed intravenous transplantation into recipient NSG mice. Cells were labeled with a bioluminescence reporter and treated with SMYD2i prior to the inoculation, and recipient animals received SMYD2i for the duration of the experiment. BLI and histological analysis showed significantly diminished metastatic colonization in animals receiving SMYD2i compared to vehicle-treated mice (Fig. 6e-f).

Altogether, these data validated SMYD2 as a driver of breast cancer metastases and supported the molecular mechanisms of action of the SMYD2-BCAR3-FMNLs signaling in the regulation of cell motility and invasion (Fig. 6g). Importantly, our *in vivo* analyses concurred that pharmacological inhibition of SMYD2 enzymatic activity constitutes a clinically targetable vulnerability to prevent breast cancer metastatic burden.

## DISCUSSION

Metastasis remains the major cause of breast cancer morbidity and mortality, however underlying mechanisms remain poorly understood and therapeutic strategies to prevent metastatic spread are limited^2, 46^. Our study provides key insights into the mechanisms that confer breast cancer cells the competence to metastasize. Altogether, our work uncovers the first known non-histone protein methylation signaling mechanism that regulates directly cell motility and invasion, deciphers the pro-metastatic role and mechanisms of action of SMYD2 methyltransferase in breast cancer and validates SMYD2 as a clinically actionable target to prevent metastases, the leading cause of breast cancer mortality.

Specifically, we found by single-cell RNA seq analysis that the lysine methyltransferase SMYD2, which elevated expression correlates with metastatic breast cancer and poor survival, is linked with high metastasis potential of neoplastic cells. Remarkably, we observed that SMYD2 depletion has no impact on cell proliferation and primary tumor growth as suggested in previous *in vitro* studies^13, 15^, but abrogates metastatic dissemination in mouse models of mammary tumorigenesis and human breast cancer models. These conflictual observations reflect the discrepancy resulting from *in vitro* and *in vivo* studies, illustrating the importance of more complex cellular environment such as animal models.

We and others have previously shown that SMYD2 lacks physiologic methyltransferase activity on histones. Since none of the known putative SMYD2 substrates are linked to pro-metastatic functions, we speculated that methylation of a yet uncharacterized SMYD2 target facilitates the observed phenotype. In an unbiased proteomic screen, we identified BCAR3 as a new substrate of SMYD2, monomethylated at lysine 334 (BCAR3 K334me1). We further validated SMYD2 as the principal enzyme tasked with generating physiologic BCAR3 K334me1, based on both depletion and reconstitution experiments in multiple independent breast cancer cell lines. Consistent with previously characterized tumor-promoting properties of BCAR3, particularly BCAR3 critical role as an activator of p130Cas/BCAR1-dependent cell motility^19, 36^, we observed a pivotal function for K334me1 of BCAR3 in controlling breast cancer migration and metastasis. However, our data suggests that the methylation of BCAR3 induced by SMYD2 overexpression highjacks the BCAR3-p130Cas regulatory mechanism of cell adhesion and cell protrusion by directly recruiting major actors of lamellipodia maturation, enhancing cancer cells capacity.

The chemical change introduced by methylation may alter protein catalytic activity or interaction with an effector protein that contains a methyl-binding domain. Because BCAR3 does not exhibit an enzymatic activity, we reasoned that K334 methylation may serve as an anchoring point for specific methyl-binding readers. To identify in an unbiased manner a methyl-binding event, we utilized stable isotope labeling (SILAC) coupled with quantitative mass spectrometry to perform methylation-specific pulldown from whole cell extracts. We discovered that the three proteins constituting the Formin-like protein family (FMNLs) are specific binders of K334 methylated BCAR3. FMNLs are essential in forming protrusive actin structures such as lamellipodia during cell migration and are associated with malignant cancer progression^20, 22, 38^. Those findings were unexpected because FMNLs lack any characterized methyl-binding domain such as PHD or Chromo domains^47^. Our analysis suggests that the GBD-FH3 domain of FMNLs is sufficient for the interaction, and is organized as an Armadillo-repeat fold sharing structural redundancy with HEAT repeats known for selectively binding monomethylated H4K20^40^. Therefore, our data suggests that we identified a novel methyllysine reader domain within a family of cytoplasmic proteins, further emphasizing the importance of lysine methylation signaling beyond chromatin and epigenetics. Interestingly, while Formins activity and specificity to modulate particular cell protrusions is well characterized, it remains unclear how each Formin are targeted to specific protrusion structures. One possibility raised by our study is that these proteins are recruited and locked to specific nascent cell protrusions based on their capacity to bind anchoring methylated proteins such as BCAR3.

Identification of BCAR3 methylation-specific interaction with FMNLs suggests that BCAR3 acts as an adapter protein coupling FMNLs effectors to p130Cas/BCAR1, a sensor of focal cell adhesion and environmental stimuli, promoting the localized formation of lamellipodia. The question remains, however, as to how FMNLs are activated upon binding to methylated BCAR3. A potential indication is provided by the mechanisms of FMNLs activation by Cdc42 GTPase^20, 22, 41^. Cdc42 binds to FMNLs via the GBD-FH3 domain and triggers FMNLs localization and activation at lamellipodia. Cdc42-mediated activation displaces the autoregulatory DAD domain (Diaphanous-autoregulatory domain) from the GBD-FH3 domain, rendering the FH1 and FH2 domains accessible for actin filament polymerization. Interestingly, our structural model suggests that methylated BCAR3 binds to the GBD-FH3 domain and may also trigger conformational changes that activate FMNLs. Alternatively, BCAR3 methylation recruits FMNLs to the leading edge of lamellipodia, where other factors may activate FMNLs. Indeed, while the potential GEF activity of BCAR3 has been ruled out^48^, several lines of evidence have shown that the interaction of BCAR3 with BCAR1 triggers downstream pathways culminating in the activation of Cdc42-GTP^49^. This would provide a crucial feed-forward mechanism, where BCAR3 methylation controls both localization and activation of FMNLs. It suggests an intriguing possibility where BCAR3 methylation by SMYD2 and FMNLs activation are regulated in a temporal- and spatial-specific pattern. Interestingly, BCAR3 is also regulated by phosphorylation at Y117, which elicits ubiquitin-dependent degradation and low BCAR3 levels in mammary epithelial cells that impede epithelial migration^50^. Hence, it is tempting to speculate that BCAR3 levels and functions are specifically regulated by different PTMs in normal tissue and in malignancy.

Our study links for the first time SMYD2, BCAR3 and FMNL proteins into a common pathway regulating cellular motility and breast cancer neoplastic cells capacity to metastasize. We discovered that SMYD2 methylates BCAR3 at lysine K334 and that this methylation mark is specifically recognized by FMNL proteins. The methylation-triggered assembly of FMNLs facilitates actin filament elongation at the cell leading edge, which generates force enabling cellular motility critically important for metastatic dissemination (Fig. 6g). Finally, our study provides the rationale for therapeutic targeting of SMYD2 activity to prevent breast cancer cells invasiveness and to impede metastasis. Specifically, we showed that the SMYD2-BCAR3- FMNL signaling pathway could be effectively disrupted by inhibiting SMYD2 enzymatic activity both *in vitro* and *in vivo*. Additionally, our data indicated that SMYD2 inhibitors are well tolerated in pre-clinical animal models and are promising therapeutics to prevent breast cancer metastatic spread, a major unmet clinical need for breast cancer patients.

## MATERIALS and METHODS

### Bioinformatic analyses

We analyzed publicly available single-cell RNA sequencing datasets of breast cancer patients (GEO; accession number: GSE176078)^12^ consisting of 130,246 cells derived from 26 primary tumors including 11 ER+, 5 HER2+ and 10 TNBCs, from which 24,489 malignant epithelial cells were selected to assess their metastasis potential. Metastasis signatures were extracted from a previous study^23^ and assigned to each cancer cell using UCell^51^. The cells were ranked according to the metastasis signature scores and were further categorized into a “Metastasis- potential-high” group (top 25%) and a “Metastasis-potential-low” group (bottom 25%). Differentially expressed genes between the two groups were identified by FindMarkers function in Seurat^52^. The Log_2_ fold change of the average expression of active lysine methyltransferases was then scaled to calculate the Z-score.

For survival analysis, we used RNA-seq data of breast cancer samples available from the public repository of The Cancer Genome Atlas (TCGA-BRCA cohort, n = 1212 samples). Briefly, we downloaded the RNA-seq SEM raw values directly provided by the TCGA firebrowse pipeline. The RSEM raw values were then transformed by DESeq. Several bio-clinical data are also available, including information about metastasis and disease-free/overall survival data. The number of events (relapses or deaths) is 199, corresponding to 16.5% of the patients. All statistical tests were performed with R software. We introduced an expression threshold for the *SMYD2* and *EZH2* genes and considered two groups of patients, those for whom the expression level of the gene in the tumor was below the threshold (group of “low” expression) and those for whom it was above the threshold (group of “high” expression). The threshold was set up to the median expression value. We then compared overall survival probabilities between the two groups of patients with “low” or “high” expressions using the log-rank statistical test. We considered the association with survival significant if the obtained log-rank p-value was less than 0.05. The survival analysis was performed using the “survminer” package of R.

Other gene expression studies were performed using TNMplot^53^ and bc-GenExMiner v4.8, compiling TCGA and SCAN-B RNA Seq datasets^54^.

### Animal models

*MMTV-PyMT, MMTV-Cre, Smyd2^LoxP/LoxP^* mice have been described before (PMID: 1312220, 9336464, 21677750, 26988419). *Smyd2^LoxP/LoxP^*mice were backcrossed to FVB/N strain for 6 generations. *MMTV-PyMT, MMTV-Cre* mice were maintained on an FVB/N strain background. and we systematically used littermates as controls in all the experiments. Immunocompromised female NSG mice (*NOD.SCID-IL2Rg-/-*) mice were used for transplantation studies. All experiments were performed on 6 to 10-week-old female animals. All animals were numbered and experiments were conducted in a blinded fashion. After data collection, genotypes were revealed and animals were assigned to groups for analysis. For treatment experiments mice were randomized. None of the mice with the appropriate genotype were excluded from this study or used in any other experiments. All mice were co-housed with littermates (2–5 per cage) in the pathogen-free facility with a standard controlled temperature of 72 °F, with a humidity of 30–70%, and a light cycle of 12 h on/12 h off set from 7 am to 7 pm and with unrestricted access to standard food and water under the supervision of veterinarians, in an AALAC-accredited animal facility at the University of Texas M.D. Anderson Cancer Center (MDACC). Mouse handling and care followed the NIH Guide for Care and Use of Laboratory Animals. All animal procedures followed the guidelines and were approved by the MDACC Institutional Animal Care and Use Committee (IACUC protocol 00001636, PI: Mazur). Tumor size was measured using a digital caliper and tumor volume was calculated using the formula: Volume = (*width*)^2^ × *length* / 2 where *length* represents the largest tumor diameter and *width* represents the perpendicular tumor diameter. The endpoint was defined as the time at which a progressively growing tumor reached 20 mm in its longest dimension as approved by the MDACC IACUC protocol (00001636, PI: Mazur) and in no experiments was this limit exceeded.

### Cell lines and patient-derived cancer xenografts

Cell lines were grown either in Dulbecco’s modified Eagle’s (293T, MDA-MB-157, MDA-MB-231, MDA-MB-453) or RPMI (MCF7) media supplemented with 10% fetal bovine serum and penicillin-streptomycin. All cells were cultured at 37°C in a humidified incubator with 5% CO2. All cell lines were authenticated by short tandem repeat profiling and tested negative for mycoplasma. Metastatic PDX was obtained from the NCI Patient-Derived Models Repository (PDMR), NCI-Frederick, Frederick National Laboratory for Cancer Research: patient (age 62) tumor specimen resected from lymph node metastasis with histologically confirmed TNBC, caring mutation in *Tp53^G^*^154^*\*,* previously treated with cyclophosphamide, doxorubicin, and paclitaxel. All tumor specimens were collected after written patient consent and in accordance with the institutional review board-approved protocols of the University of Texas M.D. Anderson Cancer Center (PA19-0435, PI: Mazur).

### Tumor and metastasis studies in GEM models

*MMTV-PyMT; MMTV-Cre; Smyd2^LoxP/LoxP^* and *MMTV-PyMT; MMTV-Cre* female mice were euthanized at indicated time and at the endpoint. Mammary tumor-free survival was determined by palpation. Whole mammary glands, tumors, and lung tissues were collected, weighed, and processed for histopathological analysis. Lung metastases were analyzed by gross examination of freshly dissected lungs and histopathological review of hematoxylin and eosin HE-stained lung sections. For therapy studies mice were treated as indicated with BAY598 (50 mg/kg daily, IP), or vehicle (0.5% hydroxypropyl methylcellulose).

### Experimental metastasis assays in mouse xenograft

Cancer cells were injected into the tail vein of 8-week-old female NSG mice: 2 × 10^5^ MDA-MB-231 cells or 1 × 10^6^ MCF7 cells. Mice were euthanized when they met the institutional euthanasia criteria for an overall health condition. The lungs were collected and processed for histopathological analysis. For PDX tumor metastasis resected tissues were propagated by orthotopically implanting small tumor fragments into the mammary fat pad of female NSG mice. Next, tumors were resected and dissociated into single cells using the MACS tumor cell dissociation kit according to the manufacturer protocol. The dissociated human tumor cells were then isolated from the contaminating mouse cells using the MACS mouse cell depletion kit following the manufacturer protocol. The cell viability and count were determined using Countess II automated cell counter and approximately 5 × 10^5^ cells were injected in female NSG mice tail vein to facilitate experimental metastasis. Mice were euthanized when they met the institutional euthanasia criteria for the overall health condition. For SMYD2 inhibitor therapy studies cells were pretreated with 5 µM BAY598 for 5 days before injection. Mouse models were treated with BAY-598 (50 mg/kg daily, IP), or vehicle (0.5% hydroxypropyl methylcellulose). For inducible expression models, cancer cells were pretreated with doxycycline (100ng/ml) for 5 days before injection, and mice were fed with 625 mg/kg Doxycycline hyclate diet starting at 5 days before cancer cells injection and were maintained on this diet for the remainder of the experiment.

### Bioluminescent imaging of metastasis in mouse xenograft

For *in vivo* metastasis assays with bioluminescent imaging, cancer cells were electroporated with a PiggyBac transposon plasmid expressing AkaLuc, which catalyzes the oxidation reaction of a substrate AkaLumine and produces near-infrared bioluminescence that can penetrate most animal tissues. To monitor metastatic spread, mice were injected i.p. with 3::Jμmol (1.0::Jmg) of AkaLumine-HCl in 100::Jμl 0.9% NaCl. Immediately after substrate injection, bioluminescent images were acquired in an AMI HTX bioluminescence imaging system. Imager settings were: emission filter, open; field of view, 25::Jcm; f-stop 1.2; low binning 2 × 2 and exposure time, 30::Js. X-ray imaging camera settings were: field of view, 25::Jcm; low exposure, and high resolution. Images were analyzed using Aura software and quantified in radiance units of photons per second per square centimeter per steradian (photons/s/cm^2^/sr) and plotted as mean ± s.e.m.

### Histology and immunohistochemistry

Tissue specimens were fixed in 4% buffered formalin for 24 hours and stored in 70% ethanol until paraffin embedding. 3 μm sections were stained with hematoxylin and eosin (HE) or used for immunohistochemical studies. Immunohistochemistry (IHC) was performed on formalin-fixed, paraffin-embedded mouse and human tissue sections using a biotin-avidin HRP conjugate (Vectastain ABC kit) method as described before (Mazur et al., 2014). The following antibodies were used (at the indicated dilutions): cleaved Caspase 3 (1:100), and Ki67 (1:1,000). Sections were developed with DAB substrate and counterstained with hematoxylin. Pictures were taken using a PreciPoint M8 microscope equipped with the PointView software. Analysis of the tumor area and IHC analysis was done using ImageJ software.

### Cell culture, transfections, and transductions

For transient expression cells were transfected with Lipofectamine 2000 transfection reagent and collected 36 h after transfection. For stable expression, 293T cells were transfected with lentiviral pSICOR (GFP/HA-tagged BCAR3 WT and K334A) using the packaging vectors pVSVg and pΔ8.2 and retroviral pMSCV (FLAG-tagged FMNL3) construct using packaging pVSVg and pUMCV vectors. Virus particles were then collected and filtrated and used for infection of relevant cells. For GFP-expressing cells, positive cells were sorted by Fluorescent Activating Cell Sorting (FACS). Cells expressing FMNL3 were selected with 10 µg/ml of blasticidin for two weeks. For constitutive or inducible knockdown, cells were transfected with pLKO.1 or pLKO-Tet-On vectors containing specific shRNA target sequences, using the packaging vectors pVSVg and pΔ8.2. Virus particles were then collected and filtrated and used for infection of relevant cells, followed by 2 µg/ml puromycin selection for one week. All cells were cultured at 37°C in a humidified incubator with 5% CO_2_.

### Identification of SMYD2 substrates

The list of potential SMYD2 substrates was generated based on the SILAC-3xMBT pulldown method already described ^55^. Briefly, HeLa cells were cultured in either normal isotope amino acids culture condition (‘Light”, K0, R0) or using modified isotope amino acids culture condition (‘Heavy’, K6, R8) for a minimum of 2 weeks. Cells were harvested, and filtered cytoplasmic extracts from hypotonic lysis were then dialyzed against KMT reaction buffer (50 mm Tris (pH 8.0), 20 mm KCl, 5 mm MgCl2, and 10% glycerol) using Slide-A-Lyzer MINI dialysis devices with 3.5K molecular weight cutoff (Pierce) and cleared by centrifugation at 15,000 × g for 5 min at 4 °C. 20 μg of recombinant SMYD2 or F184A mutant with N-terminal GST fusion was added to 1mg of light and heavy lysate supplemented with 100 μm AdoMet. Identical reactions were prepared in parallel, with light and heavy labels reversed. The reactions were incubated for 4 h at 37 °C and stopped by placing them on ice and adding 10 mm EDTA and 0.1% Triton-X. Each pair of light and heavy lysates was combined and incubated overnight on 30 μl of glutathione-Sepharose saturated with 3xMBT ^29^. Bound proteins were recovered, separated by SDS-PAGE, and processed by in-gel digestion with trypsin. Peptides were desalted using C18 stage tips (Thermo Scientific), separated by HPLC using an Ekspert nanoLC 420 (AB Sciex), and analyzed with an Orbitrap Elite mass spectrometer (Thermo Scientific). Acquisition used a data-dependent selection of the top 10 ions with a dynamic exclusion, followed by collision-induced dissociation or higher-energy collisional dissociation and analysis of fragment ions in the Orbitrap. Data were analyzed using MaxQuant version 1.3.0.5 ^56^ with a 1% false discovery rate for proteins and peptides and allowing as variable modifications methionine oxidation; acetylation of protein N termini; and mono-, di-, and trimethylation of lysine. Candidate methylation sites were verified by manual inspection.

### Identification of methylation site

For LC-MS/MS analysis of recombinant BCAR3 methylation, deuterated SAM was used to rule out possible artifactual chemical methylation *in vitro*, shifting the mass of one methyl group from 14.016 Da to 17.034 Da. After SDS-PAGE separation and Coomassie blue staining (GelCode Blue, Thermo) recombinant methylated BCAR3 (see methylation assay method) was sliced from the gel and digested with modified trypsin (sequencing grade, Promega). The resulting peptides were analyzed by online nano liquid chromatography (LC)-MS/MS (UltiMate 3000 RSLCnano and Q-Exactive HF, Thermo Scientific). For this, peptides were sampled on a 300 µm x 5 mm PepMap C18 precolumn (Thermo Scientific) and separated on 75 µm x 250 mm C18 columns (Reprosil-Pur 120 C18-AQ, 1.9 μm, Dr. Maisch). MS and MS/MS data were acquired using Xcalibur (Thermo Scientific). Mascot Distiller (Matrix Science) was used to produce mgf files before identification of peptides and proteins using Mascot (version 2.7) through concomitant searches against a database containing the sequences of proteins of interest (homemade), classical contaminants database (homemade) and the corresponding reversed databases. The Proline software ^57^ was used to filter the results (conservation of rank 1 peptides, peptide length ≥ 7 amino acids, identity threshold of peptide-spectrum-match < 0.01, minimum peptide-spectrum-match score of 25, and minimum of 1 specific peptide per identified protein group). Peptides of interest were subsequently targeted by LC-Parallel Reaction Monitoring using the same nanoLC-MS system. Candidate methylation sites were verified by manual inspection.

### Identification of methyl-sensitive binders

BCAR3 peptides were generated by Covalab based on the following sequence: Biotin-Ahx- DRRALSLKAHQSESY-CONH2. For peptide pull-down, 10 µl of Streptavidin Sepharose beads (GE Healthcare) were saturated with 7.5::Jµg of specific biotinylated peptides in peptide buffer (50 mM Tris pH 8, 150 mM NaCl, 0.5 % NP40, 0.5 mM DTT, 10 % glycerol, complete protease inhibitors (Roche)) for 2 h at 4°C under rotation. Next, beads were washed in the peptide buffer and incubated for 4 h at 4°C under rotation with 1mg of HeLa cytoplasmic extract prepared from cells cultivated in either normal isotope amino acids culture condition (‘Light”, K0, R0) or using modified isotope amino acids culture condition (‘Heavy’, K6, R8). A 2-way experiment was performed, the ‘forward’ condition combining BCAR3-K334me0 peptide with light extract and BCAR3-K334me1 peptide with heavy extract, the ‘reverse’ condition combining BCAR3-K334me1 peptide with light extract and BCAR3-K334me0 peptide with the heavy extract. Beads of each pair of peptide pulldown were then pooled together, and extracts were resuspended in Laemmli buffer. Eluted proteins were stacked on top of a 4-12% NuPAGE gel (Invitrogen). After staining with R-250 Coomassie Blue (Biorad), proteins were digested in-gel using modified trypsin (sequencing grade, Promega), as previously described ^58^. The resulting peptides were analyzed by online nano liquid chromatography coupled to MS/MS (Ultimate 3000 RSLCnano and Q-Exactive Plus, Thermo Fisher Scientific) using a 120 min gradient. For this purpose, the peptides were sampled on a pre-column (300 μm x 5 mm PepMap C18, Thermo Scientific) and separated in a 75 μm x 250 mm C18 column (Reprosil-Pur 120 C18-AQ, 1.9 μm, Dr. Maisch). The MS and MS/MS data were acquired by Xcalibur (Thermo Fisher Scientific). Peptides and proteins were identified and quantified using MaxQuant (version 1.6.2.10, Tyanova, Temu, and Cox 2016) using the Uniprot database (*Homo sapiens* reference proteome, 20180526 version) and the frequently observed contaminant database embedded in MaxQuant. Trypsin was chosen as the enzyme and 2 missed cleavages were allowed. Peptide modifications allowed during the search were: carbamidomethylation (C, fixed), acetyl (Protein N-ter, variable) and oxidation (M, variable). The minimum peptide length and the minimum number of unique peptides were respectively set to seven amino acids and one peptide. Maximum false discovery rates - calculated by employing a reverse database strategy - were set to 0.01 at peptide and protein levels. Quantification of SILAC ratios was performed using default settings. Proteins identified as outliers in both experiments are assigned as significant interactors. Amino acid complements (L-lysine-2HC, L-arginine-HCl, 2H4-L-lysine-2HCl, 13C6-L-arginine-HCl, L-proline), media, and serum used for SILAC were purchased from Silantes.

### Peptide pulldown

To prepare the resin, 5 µg of the biotinylated peptide was incubated with 4 µl streptavidin resin by gently rocking for 30 minutes at 4°C in a reaction buffer containing 20 mM Bis-Tris pH 6.5, 150 mM NaCl and 0.05% NP-40. Then the resin was washed 2 times in the same buffer. For peptide pulldown with recombinant proteins, 2 µg of protein was diluted in 200 μl of reaction buffer and directly added to the washed peptide-bound resin. This solution was incubated for 3 h rotating at 4°C. The supernatant was removed, and the beads were washed 5 times with 1 ml of reaction buffer. After removal of the final wash, the beads were resuspended in 50 μl of 2× SDS loading buffer (45 mM Tris pH 6.8, 10% glycerol, 1% SDS, 50 mM DTT, 0.002% bromophenol blue). The samples were incubated at 95°C for 10 min and 5-10 μl of the sample was resolved with an 8-12% polyacrylamide gel. For peptide pulldown with protein from mammalian protein extraction, cells were first disrupted by gently rocking for 20 minutes at 4°C in lysis buffer containing 50mM Tris pH 8.0, 500 mM NaCl, 1% Triton X-100, 1 mM EDTA, 1 mM EGTA and freshly supplemented with 1 mM DTT, 0.5 mM PMSF and anti-proteases. Then cell debris was discarded by centrifugation, and supernatant was collected. For each pulldown experiment, 500 µg to 1 mg of total protein was loaded on the washed peptide-bound resin. This solution was incubated for 16 h rotating at 4°C. The supernatant was removed, and the beads were washed 4 times with 500 μl of reaction buffer. After removal of the final wash, the beads were resuspended in 50 μl of 2× SDS loading buffer. The samples were incubated at 95°C for 10 min, and 10-15 μl of the sample was resolved on an 8-12% polyacrylamide gel.

### Expression and purification of recombinant proteins

BCAR3 and SMYD2 recombinant proteins were purified from transformed BL21 bacteria cells, induced with 0.1 mM IPTG for 16 h at 16°C. Cells were resuspended in lysis buffer containing 50 mM Tris pH 7.5, 150 mM NaCl, 0.25 mg/ml lysozyme, 0.5 mM PMSF and protease inhibitors, and additionally lysed by sonication. Full-length FMNL1, 2 and 3 were purified from transformed BL21 Arctic strain bacteria cells, induced with 0.25 mM IPTG for 36 h at 10°C. Cells were resuspended in high salt lysis buffer containing 50 mM Tris pH 8.0, 500 mM NaCl, 10% glycerol, 0.25 mg/ml lysozyme, 0.5 mM PMSF, 1 mM DTT and protease inhibitors, and additionally lysed by sonication. The fragment FMNL3 (1-382) was purified from transformed BL21 Rosetta-2, induced with 0.25mM IPTG for 16h at 16°C. Cells were resuspended in lysis buffer containing 100 mM Tricine pH 8.0, 500 mM NaCl, 10% glycerol, 0.25 mg/ml lysozyme, 0.5 mM PMSF, 1 mM DTT and protease inhibitors. GST-tagged proteins were purified using Glutathione Sepharose 4B beads and eluted in 10::JmM reduced L-glutathione in 100mM Tris pH 8.0, or 100mM Tricine pH 8.0 for FMNL3 (1-382).

### Methylation assay

*In vitro* methylation assays were performed by using 2 µg of recombinant proteins which were incubated overnight with 2 µg of SMYD2 and either 0.1 mM S-adenosyl-methionine (SAM) or 0.1 mM S-adenosyl-l-methionine-d3 tetra (p-toluenesulfonate) salt (deuterated SAM, CDN isotope) or 2::JµCi SAM[^3^H] (IsoBio) in buffer containing 50 mM Tris-HCl (pH 8.0), 10% glycerol, 10 mM KCl, 5 mM MgCl_2_ at 30°C overnight. In the case where SMYD2 inhibitor (BAY-598) was used, the enzyme was preincubated for 1h in the buffer at 4°C with the inhibitor or the vehicle before adding substrate. The reaction mixture was resolved by SDS-PAGE, followed by autoradiography, western blotting, Coomassie stain, or mass spectrometry analysis.

### Immunoprecipitation and co-immunoprecipitation

For immunoprecipitation and detection of endogenous BCAR3 methylation, cells were cultivated 48h to 72 h with either 100 ng/ml of doxycycline or 5 µM of BAY-598 (SMYD2i) before harvesting. The cells were resuspended in lysis buffer (50 mM Tris-HCl pH 8.0, 500 mM NaCl, 1% Triton X-100, 1 mM EDTA, 1 mM EGTA and freshly supplemented with 1 mM DTT, 0.5 mM PMSF and anti-proteases. Lysates were then incubated with magnetic beads coupled to protein-A, containing either BCAR3 antibody or BCAR3 K334me1 antibody. After 16h rotating at 4°C, beads were washed 3 times and proteins were eluted in 2× SDS loading buffer. For immunoprecipitation of ectopic proteins, immunoprecipitation of HA-tagged BCAR3 WT or K334A was completed after transient expression in 293T cells for 36 h. Cells were lysed as before and incubated in pre-washed anti-HA resin. After 16h rotating at 4°C, beads were washed 3 times and proteins were eluted in a 2× SDS loading buffer. For the co-immunoprecipitation of FMNL2/3 with HA-GFP BCAR3 in engineered MDA-MB 231, Tris-HCl was replaced with 50mM Tricine pH 7.8. The cell lysate was cleared by centrifugation, the buffer was adjusted to 150mM NaCl and 0.1% Triton and soluble proteins were added to pre-washed anti-HA resin. Similarly, for co-immunoprecipitation of FLAG-FMNL3 with HA-GFP BCAR3 in engineered MDA-MB 231, the lysate was added to prewashed anti-FLAG resin. For endogenous co-immunoprecipitation of FMNL2/3 and BCAR3, the lysate was prepared as before and added on agarose resin coupled to protein A, incubated with FMNL2/3 antibody beforehand. After overnight incubation at 4 °C with rotation, HA-resin, FLAG resin, or FMNL2/3- protein A resin with bound proteins were washed three times in the same adjusted buffer. Proteins were eluted from the beads with a 2× SDS loading buffer and analyzed by western blot.

### Immunoblot analysis

For western blot analysis, cells were lysed in RIPA buffer with 1 mM PMSF and protease inhibitor cocktail. Protein concentration was determined using the Coomassie plus assay. Protein samples were resolved by SDS-PAGE and transferred to a PVDF membrane. The following antibodies were used (at the indicated dilutions): SMYD2 (1:1000), BCAR3 (human, 1:1000), BCAR3 (mouse, 1:1000), BCAR3 K334me1 (1:1000), FMNL1 (1:1000), FMNL2 (1:1000), FLAG (1:1000), HA (1:1000), Tubulin (1:2000), Biotin (1:5000). Secondary antibodies were used at 1:5000 dilution. Protein bands were visualized using an ECL detection reagent.

### Homology modeling of FMNL3 GBD-FH3 domain

The structural model of the FMNL3 GBD-FH3 domain was built by using the SWISS-MODEL server ^60^ by adding as templates the X-ray structures of human FMNL2 GBD-FH3 / Cdc42 GppNHp (PDBe code: 4YC7); and the structure of human FMNL1 (PDB code: 4YDH). The sequence identities between FMNL3 GBD-FH3 (residues 33 to 419) and FMNL1 and FMNL2 were 80.0 and 88.6% respectively. The calculated homology model of the FMNL3 GBD-FH3 domain presented Qmean Z scores of -2.86 and 3.14, indicating a very good global-quality model ^61^. The root-mean square deviations calculated in Coot ^62^ over carbon-alpha atoms of FMNL3 GBD-FH3 with respect to FMNL1 and FMNL2 were 0.18 and 0.74 Å respectively. The PyMOL Molecular Graphics System (v.2.5.1, Schrodinger, LLC) was used to prepare structural models.

### In silico docking of methylated K334-BCAR3 into FMNL3 GBD-FH3 domain

The secondary structure propensity of BCAR3 residues 319 to 397 was calculated using the prediction sever PSIPred ^63^, indicating that the region with residues 321 to 342, bearing Lysine 334, is predicted to fold into a helix. The three-dimensional structure of this BCAR3 helix and its docking into FMNL3 were calculated in Coot ^62^ via mutagenesis and local energy refinements by using as a template model the structure of Cdc42 in complex with FMNL2 (PDB code : 4YC7). Specifically, the position of methylated K334 BCAR3 was modeled by using the guanidinium group of Arginine 65 within helix 3 of Cdc42. No steric clashes or violations of the interatomic Van der Waals radii were found after BCAR3 K334me1 was placed at the hydrophobic pocket of FMNL3 GBD-FH3.

### Live microscopy analysis of migration and protrusions

For collective migration with wound healing analysis, cells were seeded on Ibidi culture inserts and allowed to grow for 48-72 h until cells formed a homogenous monolayer. Then, the insert was carefully removed, and cell monolayers were washed 4 times with complete DMEM. Imaging was performed immediately away after selecting 1 or 2 areas per condition, with 1 image every 15 minutes for 24 h. Wound closure was calculated as the ratio of the area covered by the cells over the remaining empty area at each time point. For random cell migration, cells were seeded on a 4-well chamber Ibidi coated pH+ and allowed to grow for 48-72 h with or without doxycycline (100ng/ml). Brightfield images were acquired with 10× objective at 37::J°C with 5% CO2 for 15::Jh every 5::Jminutes. Only cells that did not undergo mitosis for 12h were considered. Velocity and migration patterns of individual cells were tracked manually employing the built-in Fiji plugin Manual Tracking. Analysis of migration speed was conducted using the program developed by ^64^. For protrusion analysis, cells were seeded on a 4-well chamber Ibidi coated pH+ and allowed to grow for 48-72 h with or without doxycycline (100ng/ml). Brightfield images were acquired with a 33x objective at 37::J°C with 5% CO2 for 15 minutes every 3 seconds. Kymographs were then generated using the built-in Fiji plugin and protrusion speed was calculated using the slope manually drawn for each lamellipodia over 10 minutes.

### Spheroid invasion assay and adaptive optics

To perform the analysis 2×10^3^ cells were seeded in a 96-well plate with a ULA round bottom, centrifugated at 300 g and allowed to grow for 24 h. Then basement membrane Matrigel was added at a concentration of 250 µg/ml, and cells were centrifugated at 300 g and let in culture for 36 h until they formed compact spheroid. Each spheroid was then carefully transferred into cold and liquid collagen I mixture at 1.6 mg/ml, cast in 4-well Ibidi. Gels containing spheroids were allowed to polymerize for 2-3 h at 37°C in a humified incubator. First brightfield images (t = 0 h), and last ones (48 h) of each spheroid were taken with a 10× objective in a homemade microscope chamber, maintaining cell culture condition. Those images were used to quantify the spheroid radial area. To measure the area of invasion, the total area covered by each spheroid at t = 48h was subtracted from the initial area captured by the images acquired 3 h after spheroid embedding and referred to as t = 0 h. At 48 h, gels containing spheroid were fixed in 4% PFA with 7% sucrose for 2 h at 37°C. After extensive washing, spheroids were permeabilized with PBS containing 0.2 % Triton for 30 minutes, washed 5 times in PBS and incubated with Texas-Red Phalloidin (1: 250) and Hoescht (1: 200) for 16 h at 4°C. Gels were finally washed 5 times in PBS, and imaged right away. Due to their volume and the divergence index of the gel, spheroids were challenging to image without losing focus. To circumvent this issue, we employed adaptive optics to correct optical aberrations and then get image acquisition of the cells deeper within the gel. Those images were chosen to illustrate spheroid morphology upon different conditions tested.

### Proximity Ligation Assay (PLA)

Cells were cultivated on coated coverslips with 20 µg/ml fibronectin, washed two times with PBS and fixed in cold methanol at -20 °C for 5 minutes. Fixed cells were blocked with Duolink Blocking Solution for 2::Jh. Primary antibodies (rabbit BCAR3 K334me1, mouse HA, rabbit FLAG) were added overnight at 4::J°C. PLA reactions were subsequently carried out using Duolink PLA plus and minus probes for rabbit and mouse (respectively) and Duolink In Situ Detection Reagents Orange (Sigma) following the manufacturer’s protocol. Coverslips were finally mounted with Mowiol DAPI. Images for methylation detection were acquired with epifluorescence microscopy with a 20x objective. Images for FMNL3/BCAR3 interaction were acquired with confocal spinning disk microscopy with a 40x objective. The number of dots per cell was calculated as the ratio between the total number of dot and cell nuclei within one frame. Each frame is then represented as a point in the quantification graph.

### Immunofluorescence and confocal microscopy

Cells were cultivated on coated coverslips with 20 µg/ml fibronectin, washed two times with warm PBS, and fixed in 4% PFA - 5% Sucrose at 37°C for 15 minutes. Fixed cells were washed five times and permeabilized in PBS-0.1% Triton X-100 for 5 minutes. After five washes, specific sites were blocked using a solution of PBS-3% BSA for 1 h and coverslips were incubated overnight with indicated primary antibody for 16 h at 4°C. Coverslips were washed three times and incubated for 1h with a dedicated secondary antibody with Texas-Red Phalloidin. After five extensive washing in PBS and one last in distilled water, coverslips were mounted with Mowiol DAPI and sealed. For lamellipodia observations, images were taken with confocal spinning disk microscopy with a 40× objective for MCF-7 and a 63× objective for MDA-MB 231. The maximum intensity projection of phalloidin staining was generated with Fiji, and each lamellipodium was manually outlined. For each outline, the integrated density of lamellipodia fluorescence was assessed with Fiji, as well as a square near lamellipodia without signal and referred to as the background. Fluorescence intensity of lamellipodia (CTLF, for Corrected Total Lamellipodia Fluorescence) was then calculated as below: CTLF = Integrated density (lamellipodia) – [Mean intensity (background) × Area (lamellipodia)]. To analyze FLAG-FMNL3 enrichment at lamellipodia, images were taken with confocal spinning disk microscopy with a 40× objective. Maximum intensity projection of phalloidin and FLAG staining were generated with Fiji, and each cell as well as lamellipodia were manually outlined using phalloidin staining as a template. For each outline, the integrated density of cell and lamellipodia fluorescence from FLAG staining was assessed with Fiji, as well as a square near of cell without signal and referred to as the background. Enrichment of FLAG-FMNL3 in lamellipodia was calculated as below: FMNL3 (lam) = CTLF/CTCF with, CTCF = Integrated density (cell) – [Mean intensity (background) × Area (cell)].

### Quantification and statistical analysis

Please refer to figure legends for the description of sample size (n) and statistical details. All values for n are for individual mice or individual samples. Sample sizes were chosen based on statistical power calculations and previous experience with given experiments. Cell culture assays have been performed in triplicates and two independent experiments, unless stated otherwise. Differences were analyzed by log-rank, two-tailed unpaired Student’s t-test, and two-way ANOVA with Tukey’s testing for multiple comparisons using Prism 8 (GraphPad), unless stated otherwise.

## RESOURCE AVAILABILITY

### Lead Contact

- Further information and requests for resources and reagents should be directed to and will be fulfilled by the Lead Contact, Nicolas Reynoird.

### Materials Availability

- Plasmids, antibody and cell lines generated in this study will be available upon reasonable request from the lead contact upon request with a completed material transfer agreement.

### Data and Code Availability

- All gene expression data utilized in the study are publicly available.
- This study did not generate any unpublished code, software, or algorithm. All utilized codes are publicly available as of the date of publication.
- Any additional information required to reanalyze the data reported in this paper is available from the lead contact upon request.

## ACKNOWLEDGMENTS

Authors would like to thank Alexei Grichine, Mylène Pezet and Jacques Mazzega from the Optical Microscopy - Cell Imaging facility at the Institute for Advanced Biosciences, for their help in imagery analyses. N.R. was supported by the French National Research Agency in the framework of the "Investissements d’avenir” program (ANR-15-IDEX-02), the French National Cancer Institute (INCA_ 16066), la Ligue contre le cancer (Comité Saone et Loire, 2021) and Gefluc Isère 2020. P.K.M. was supported by NIH (R01CA236118, R01CA236949, CA266280), DoD PRCRP Career Development Award (CA181486), CPRIT IIRA (RP220391) and CPRIT Scholar in Cancer Research (RR160078). The proteomic experiments were partially supported by French National Research Agency under projects ProFI (Proteomics French Infrastructure, ANR-10-INBS-08) and GRAL, a program from the Chemistry Biology Health (CBH) Graduate School of University Grenoble Alpes (ANR-17-EURE-0003). S.H. was supported by NIH (K99 CA255936). A.C. is a recipient of a doctoral fellowship from FRM (Fondation pour la Recherche Medicale, ECO20170637491) and a 4th year PhD fellowship from ARC (Association pour la Recherche contre le Cancer). We thank members of the Reynoird and Mazur labs for their critical reading of the manuscript.

## CONFLICT of INTERESTS

Pawel K. Mazur is a scientific co-founder, consultant and stockholder of Amplified Medicines, Inc. and Ikena Oncology, Inc.

## AUTHOR CONTRIBUTIONS

A.G.C., P.K.M. and N.R. were responsible for the experimental design, execution and data analysis. A.G.C., G.S.R., L.B, Y.C. and N.R. performed and analyzed mass spectrometry experiments. A.G.C., G.S.R., J.V. and N.R. performed and analyzed biochemical and cell assays. X.L., E.F., S.R, and S.B. performed bioinformatics analyses. A.G.C., F.I., A.P. and N.R. performed structural modeling. S.H., P.K.M., A.M.B., N.M.F., M.C., performed and analyzed animal experiments, treatments and histological studies. S.R., Y.C., O.G. P.H., P.K.M. and N.R. supervised all specific aspects of the study. A.G.C., N.R. and P.K.M drafted the manuscript. P.K.M. and N.R. were responsible for funding acquisition, supervision of research, data interpretation and manuscript final preparation. All authors participated in reviewing and editing of the manuscript.

