## Supplemental Figures 1-6 for "Cytoskeleton remodeling induced by SMYD2 methyltransferase drives breast cancer metastasis"

### Extended Data Figure 1

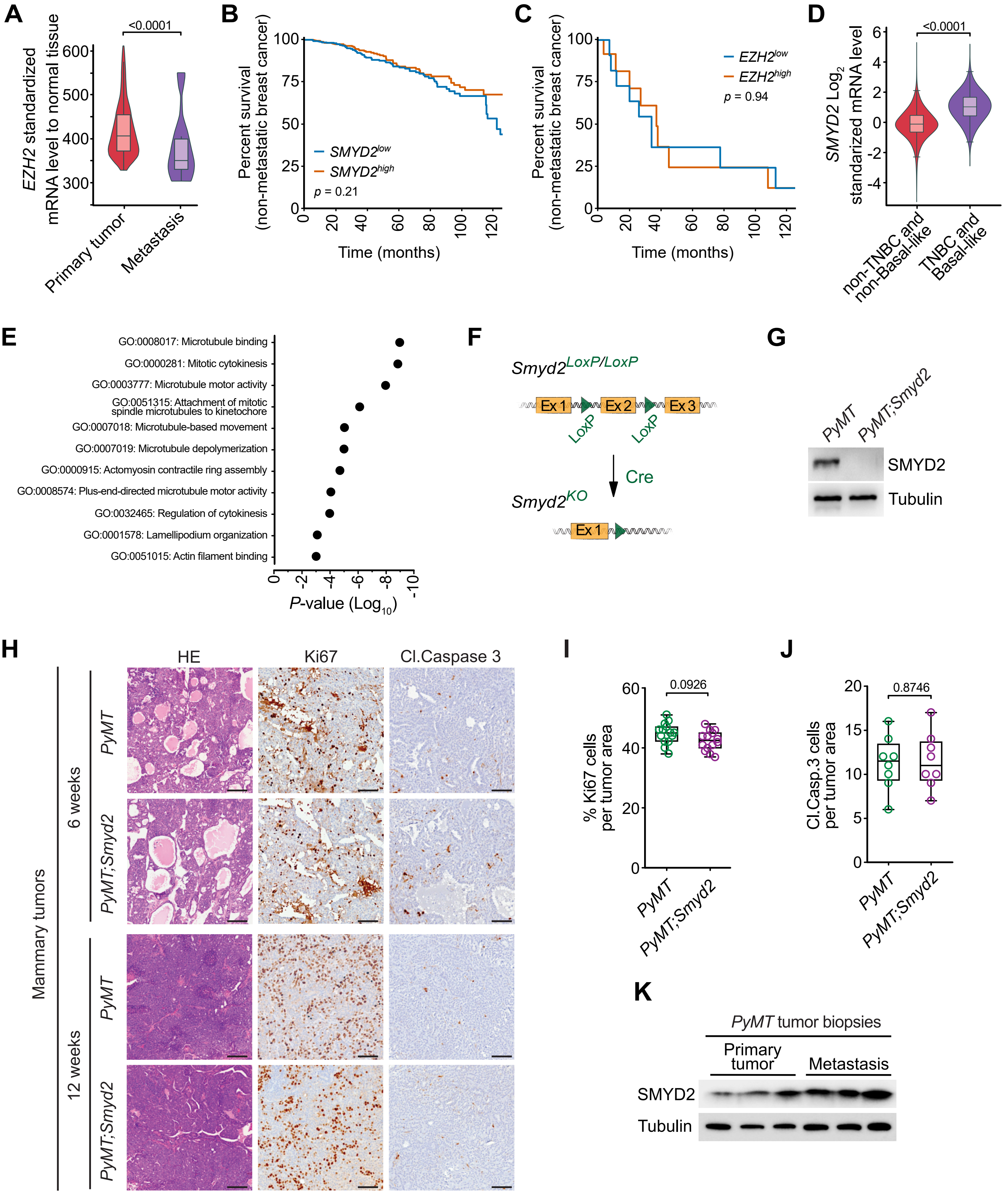

### Extended Data Figure 2

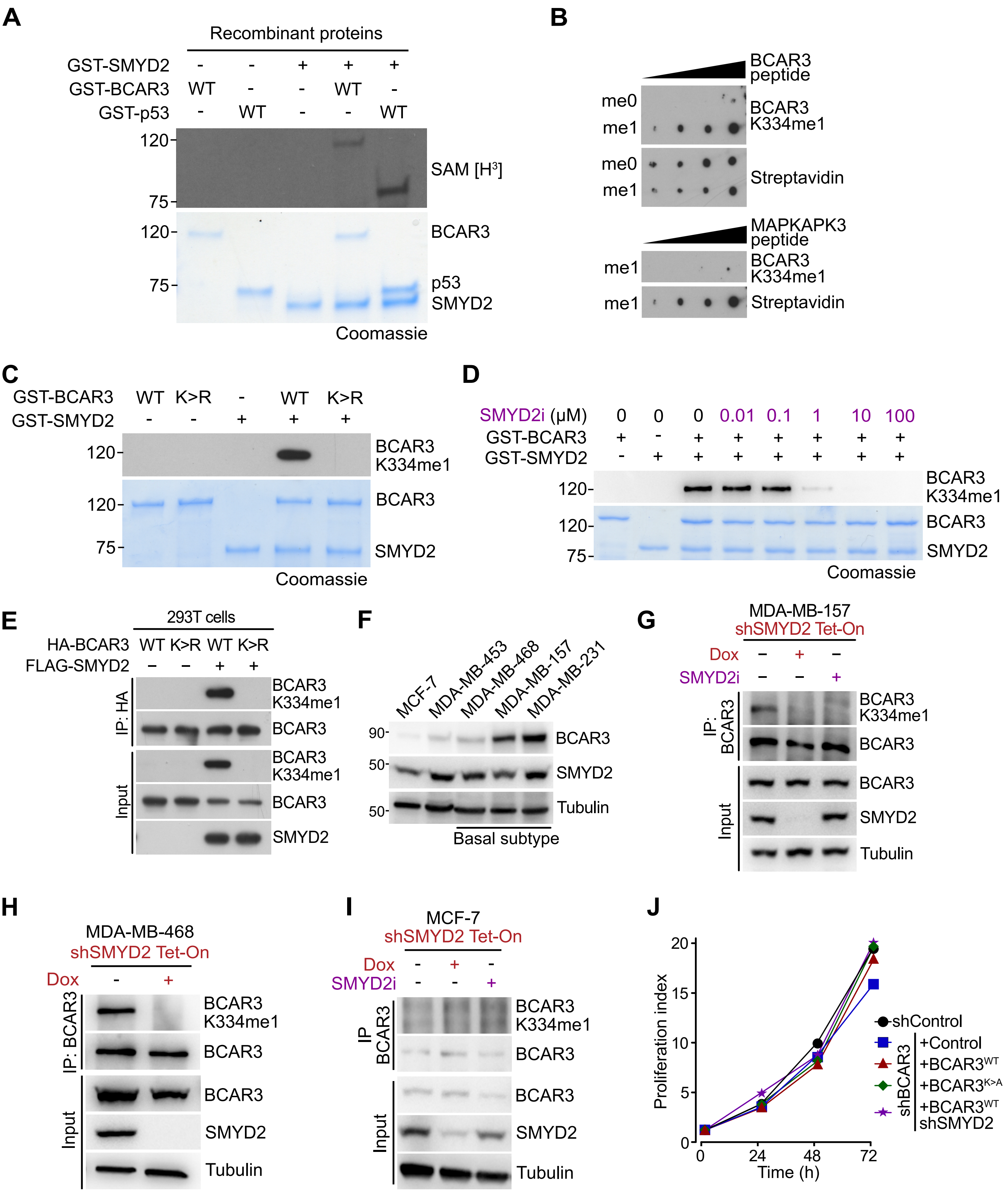

#### Extended Data Figure 3

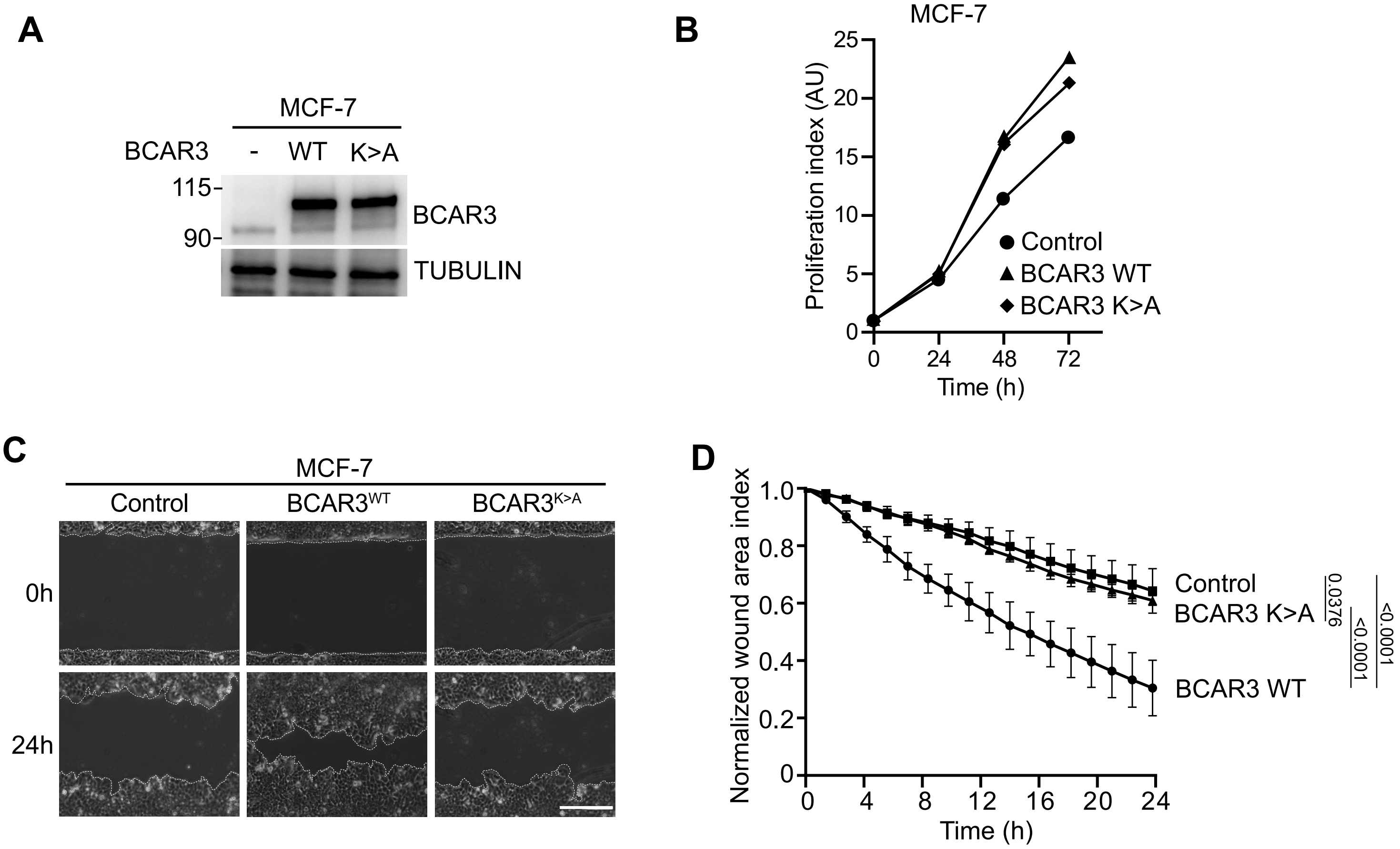

### Extended Data Figure 4

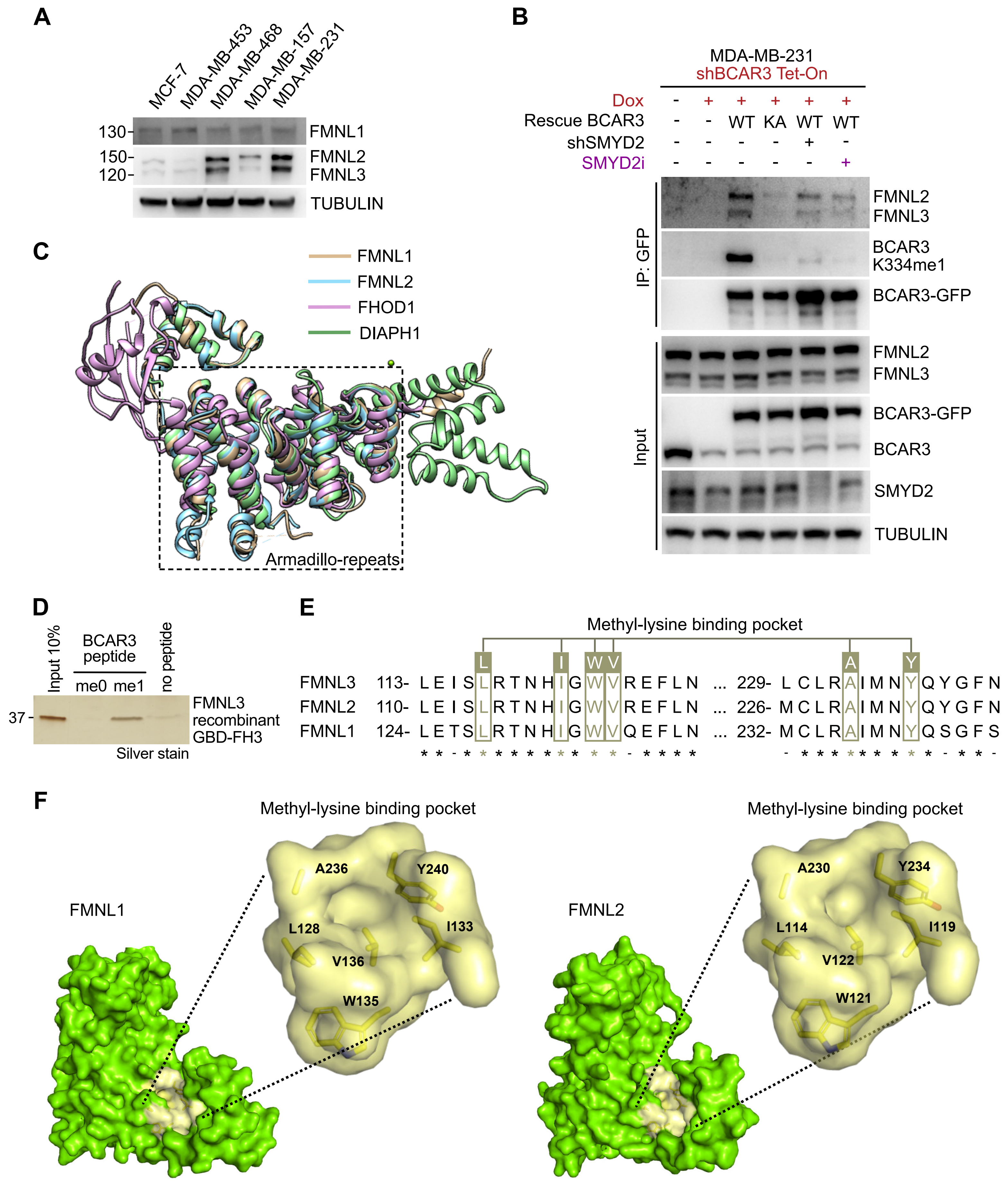

### Extended Data Figure 5

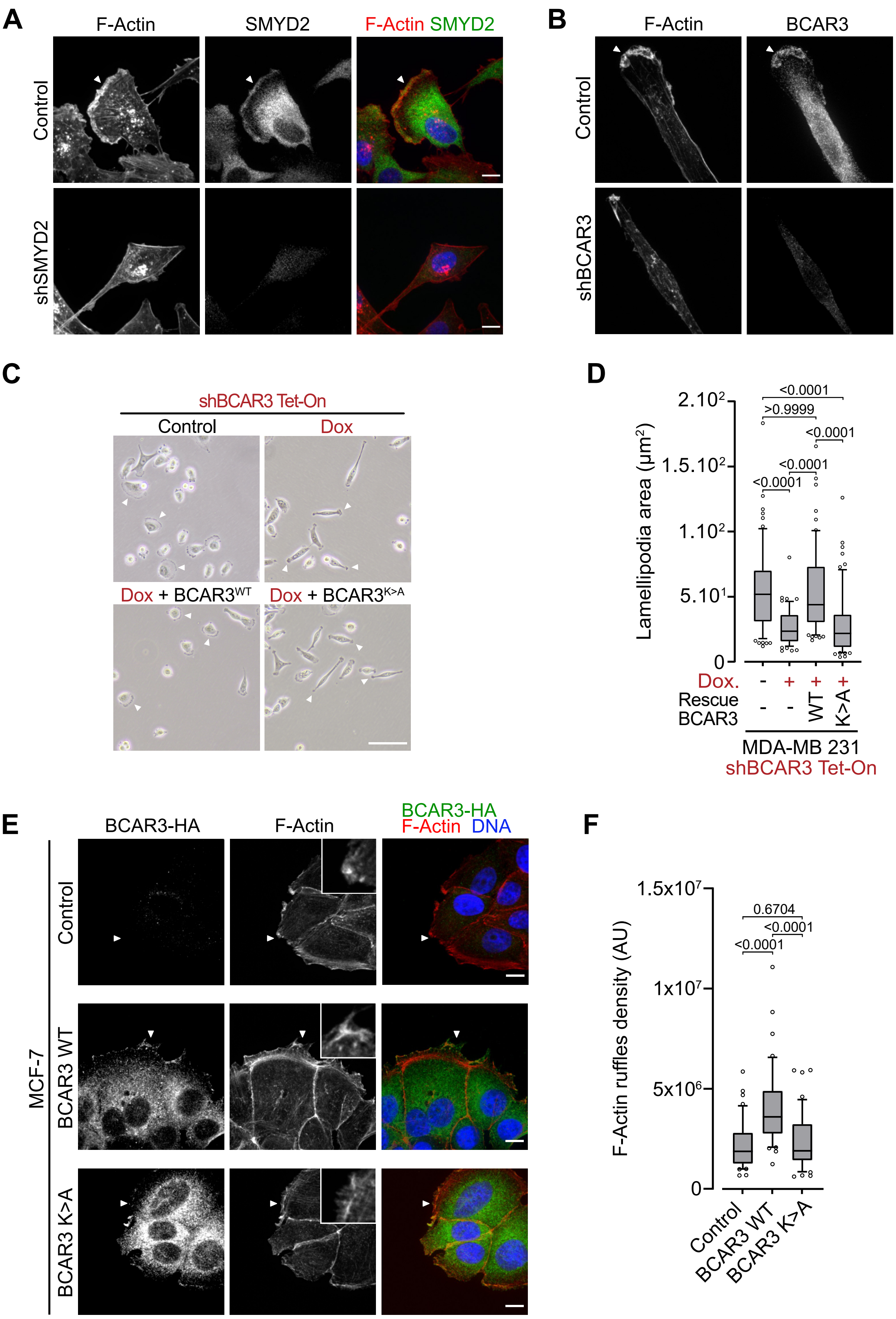

### Extended Data Figure 6

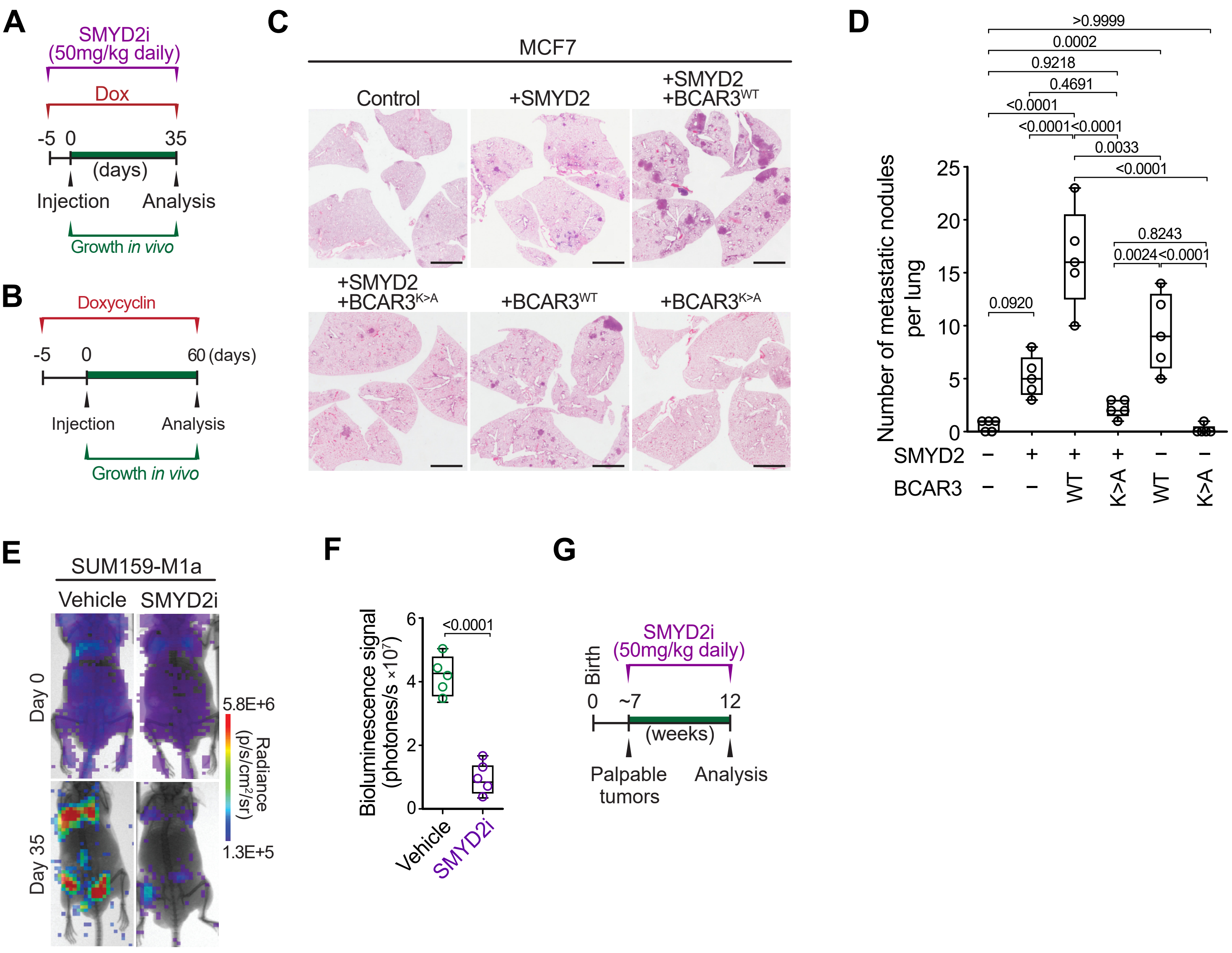
