## Supplemental information for "Cytoskeleton remodeling induced by SMYD2 methyltransferase drives breast cancer metastasis"

### SUPPLEMENTAL TABLES

#### Table S1

Gene ontology (GO) analysis identifies significantly over-represented terms in the list of genes most positively correlated with high SMYD2 expression in TNBC patient samples. Genes with Pearson's correlation coefficient above 0.40 in absolute value and associated P-value less than 0.05 were analyzed; GO analyses *P*-value calculated by Fisher's exact test; in blue, GO terms presented in Figure S1C.

#### Table S2

List of proteins identified by mass spectrometry following enrichment of methylated proteins by 3XMBT pulldowns after SILAC labeling.

#### Table S3

Spearman correlation analysis between SMYD2 and SMYD2 substrates expression levels in non-metastatic *vs* metastatic breast cancers.

#### Table S4

List of proteins identified by mass spectrometry after enrichment of labeled proteins interacting with BCAR3 K334 or K334me1 peptides.

### EXTENDED DATA FIGURE LEGENDS

#### Extended Data Figure 1. SMYD2 promotes breast cancer metastasis, related to Fig.1

**a**, Violin plots of EZH2 expression levels in primary tumor and metastases samples from breast invasive carcinoma RNA-Seq data analyzed with TNMplot. *P*-values was calculated by Kruskal-Wallis test with Dunn test for multiple comparisons. **b**-**c**, Analyses of the correlation between SMYD2 expression levels and survival in cohort of patients diagnosed with non-metastatic breast cancer (**b**) and between EZH2 expression levels and survival in cohort of patients diagnosed with metastatic breast cancer (**c**) from the TCGA RNAseq dataset. The low and high groups were set to the median expression of SMYD2 and EZH2. *P*-value were calculated by log-rank test. **d**, Violin plots showing SMYD2 expression levels in breast cancer subtypes based on TNBC/Basal-like status. Analysis was performed using bc-GenExMiner v4.8^2^. The dataset (n = 4421 samples) was defined in subgroups using PAM50 and IHC classification in non TNBC/Basal-like (n = 3479) and TNBC/Basal-like (n = 189) subgroups. *P*-values were calculated by Welch's test with Dunnett-Tukey-Kramer's test for multiple comparisons, error bars represent mean ± SD. **e**, Gene ontology enrichment analysis of genes positively correlated with SMYD2 expression in TNBC. Selected pathways are depicted and the complete analysis is shown in Table S1. Genes with Spearman correlation with coefficient greater than 0.40 and *P*-value less than 0.05 were analyzed (expression data sets: TCGA, SCAN-B: GSE96058 and GSE81538, n = 246 patient samples. **f**, Schematic of the *Smyd2* conditional allele. In the presence of Cre recombinase, exon 2 is deleted to disrupt *Smyd2* expression. **g**, Immunoblot analysis with the indicated antibodies of tumor biopsy lysates from *PyMT* and *PyMT;Smyd2* mice. Tubulin is shown as a loading control. **h**, Representative HE-stained sections and IHC staining for markers of cell proliferation (Ki67) and apoptosis (cleaved Caspase 3) of the mammary tumors of *PyMT* and *PyMT;Smyd2* mutant mice at 6 and 12 weeks of age (representative of n = 8 mice for each experimental group). Scale bars, 50 μm. **i**-**j**, Quantification of Ki67, a marker of proliferation (**i**) and cleaved Caspase 3, a marker of apoptosis (**j**) positive cells in the mammary tumors of *PyMT* and *PyMT;Smyd2* mutant mice at 12 weeks of age. Boxes: 25^th^ to 75^th^ percentile, whiskers: min. to max., center line: median. *P*-value were calculated by two-tailed unpaired t test. **k**, Immunoblots with the indicated antibodies of primary tumors or metastases biopsy lysates from *PyMT* mice (representative of n = 3 mice for each experimental group). Tubulin is shown as a loading control.). *P-*values were calculated by Fisher's exact distribution test.

#### Extended Data Figure 2. SMYD2 methylates BCAR3 in breast cancer cells, related to Fig.2

**a**, *In vitro* methylation assays with recombinant SMYD2 (GST-SMYD2) enzyme and recombinant BRAC3 (GST-BCAR3) or positive control (GST-p53) as substrates. Top panel, [^3^H] S-adenosyl methionine (SAM[H^3^]) is the methyl donor and methylation visualized by autoradiography. Bottom panel, Coomassie stain of proteins in the reaction. **b**, Specific recognition of BCAR3 K334me1 peptides by the anti- BCAR3 K334me1 antibody using dot blot analysis on the indicated biotinylated peptides. Streptavidin is shown as a loading control. **c**, Immunodetection of BCAR3 methylation by SMYD2 after non radiolabeled methylation assay using recombinant SMYD2 and recombinant WT or K334A (K>A) BCAR3. Bottom panel, Coomassie stain of proteins in the reaction. **d**, Immunodetection of BCAR3 methylation as in (**c**), with pharmacologic inhibition of SMYD2 using indicated concentrations of BAY-598 compound. Bottom panel, Coomassie stain of proteins in the reaction. **e**, Immunodetection of BCAR3 K334me1 in 293T cell lysate after immunoprecipitation (IP) of ectopically expressed wildtype (WT) or K334R (K>R) BCAR3 in presence or absence of SMYD2. **f**, Immunodetection of BCAR3 and SMYD2 in a panel of breast cancer cell lines. Tubulin is shown as a loading control. **g-i**, Immunoblot analysis with indicated antibodies of BCAR3 or BCAR3 K334me1 immunoprecipitates from MDA-MB-157 (**g**), MDA-MB-468 (**h**) and MCF7 (**i**) cell lines. Tubulin is shown as a loading control. **j**, Analysis of the proliferation of MDA-MB-231 cells with indicated engineering, related to Figure 2f.

#### Extended Data Figure 3. BCAR3 methylation promotes cell migration and invasiveness, related to Fig.3

**a**, Immunoblot analysis with the indicated antibodies of cell extracts isolate from MCF-7 cells expressing WT or K334A (K>A) BCAR3 with GFP-HA-tag. Tubulin is shown as a loading control. **b**, Proliferation index of engineered MCF-7 cells as in (**a**). **c**-**d**, Representative image of MCF-7 cell as in (**a**) in migration assay (**c**) and quantification of cell migration (**d**). *P-*values were calculated by ANOVA with Tukey’s testing for multiple comparisons, error bars represent mean ± SD. Scale bars represent 100µm.

#### Extended Data Figure 4. FMNLs are BCAR3 methyl-specific interactors, related to Fig.4

**a**, Immunoblot analysis of FMNL1-3 expression in panel of breast cancer cell lines. Tubulin is shown as a loading control. **b**, Immunodetection of FMNL2/3 and BCAR3 co-immunoprecipitation after GFP-BCAR3 enrichment in engineered MDA-MB-231 cells. Tubulin is shown as a loading control. **c**, Ribbon model superposition of the FH3 armadillo-repeats domains of Formin homology proteins: FMNL1, FMNL2, FHOD1 and DIAPH1 based on available deposited structures (PDBe of FMNL1: 4ydh; FMNL2: 4yc7; FHOD1: 3dad; DIAPH1: 3eg5). **d**, Silver staining following pulldowns with unmethylated and K334-methylated BCAR3 peptides using recombinant GBD-FH3 domain of FMNL3. **e**, Sequence alignment of FMNL1, 2 and 3 showing a strict conservation of all aromatic and aliphatic residues identified in the structural model of FMNL3 hydrophobic pocket responsible for BCAR3me1 recognition (related to Figure 4G). **f**, Structural homology of the FMNL1 and 2 hydrophobic pockets depicted around key residues identified in Figures 4G and S4D for BCAR3me recognition.

#### Extended Data Figure 5. BCAR3 methylation recruits FMNLs to lamellipodia and regulates cytoskeleton protrusions, related to Fig.5

**a**, Representative immunofluorescence imaging of SMYD2 localization in lamellipodia visualized by F-Actin (phalloidin) in MDA-MB-231 cells. SMYD2 depleted (shSMYD2) cells serve as negative control. Scale bars, 5 µm. **b**, Representative immunofluorescence imaging of BCAR3 localization in lamellipodia visualized by F-Actin (phalloidin) in MDA-MB-231 cells. BCAR3 depleted (shBCAR3) cells serve as negative control. Scale bars, 5 µm., **c**, Representative bright-field imaging cells protrusions visualized by F-Actin (phalloidin) in MDA-MB-231 with Dox-inducible depletion (Dox) of endogenous BCAR3 and ectopic expression of WT or K>A BCAR3. Scale bars, 50 µm. **d**, Quantification of lamellipodia area in engineered MDA-MB-231 cells as in (**C**). *P-*values were calculated by Brown-Forsythe and Welch ANOVA tests with Dunnett's T3 testing for multiple comparisons. **e**-**f**, Representative immunofluorescence images (**e**) and signal quantification (**f**) of F-Actin protrusion (phalloidin) in MCF-7 cells with ectopic expression of WT or K>A BCAR3. *P-*values were calculated by Brown-Forsythe and Welch ANOVA tests with Dunnett's T3 testing for multiple comparisons. Scale bars, 5 µm.

In all box plots, the center line indicates the median, the box marks the 75^th^ and 25^th^ percentiles and whiskers: 10% to 90%, center line: median.

#### Extended Data Figure 6. The SMYD2-BCAR3-FMNLs axis drive breast cancer metastasis, related to Fig.6

**a**, Experimental design to assess the metastatic ability of MDA-MB-231 cells by intravenous transplantation into recipient NSG mice. MDA-MB-231 cells with depletion of endogenous SMYD2 or Dox-inducible depletion (Dox) of endogenous BCAR3 and ectopic expression of wildtype (WT) or K334A methyl-mutant (K>A) BCAR3 or control cells were injected. Animals were treated as indicated with Dox-containing diet or treated with SMYD2 inhibitor (BAY-598, SMYD2i). Cells were engineered to express bioluminescent reporter (AkaLuc) and followed for metastatic growth *in vivo* using bioluminescent imaging. **b**, Schematic of MCF-7 cell xenografts treatment protocol. **c**-**d**, Representative HE staining (**c**) and quantification (**d**) of metastatic foci in the lungs of NSG mice injected with MCF-7 cells with ectopic expression of SMYD2 and WT or K>A BCAR3. Representative of n = 5 mice for each experimental group. Scale bars, 3 mm. Boxes: 25th to 75th percentile, whiskers: min. to max., center line: median, n = 5 mice for each experimental group, *P*-values were calculated by ANOVA with Tukey’s testing for multiple comparisons. **e**-**f**, Representative bioluminescence imaging (**e**) and signal quantification (**f**) of SUM159-M1a cells at the time of intravenous transplantation (day 0) and 35 days post-injection. Animals were treated with SMYD2 inhibitor (BAY-598, SMYD2i) or vehicle (control). Representative of n = 5 mice for each experimental group. *P*-value were calculated by two-tailed unpaired t-test. **g**, Experimental design to assess the efficacy of SMYD2 inhibitor (BAY-598, SMYD2i) to attenuate metastatic spread in *PyMT* mouse model.
